# Voltage versus concentration in ion channels: Physical signatures reveal rate-limiting steps

**DOI:** 10.64898/2026.10.22.760804

**Authors:** Hannah Weckel-Dahman, Ryan Carlsen, Orion C. Harmon, Alexander D. Daum, Jessica M.J. Swanson

## Abstract

Within the field of electrophysiology, I-V curves are a primary experimental means of characterizing ion channel behavior. By directly measuring ionic currents across membranes, electrophysiological techniques quantify key properties, including conductance, ion selectivity, and voltage dependence. We demonstrate a method of linking experimentally observed ion flux with microscopic mechanistic properties of channels. Although voltage and chemical potential can provide the same net electrochemical driving force, they perturb ion transport differently: voltage modifies transition rates throughout the permeation pathway, whereas chemical potential primarily alters ion uptake probabilities. Consequently, uptake, transfer, and release respond differently to the two driving forces, producing distinct flux signatures and causing chemical potential-driven flux to plateau while voltage-driven flux continues to increase. These differences encode the rate-limiting step (RLS) in open-channel flux. We propose a protocol based on three standard I-V measurements obtained under symmetric and asymmetric ion concentrations for elucidating the RLS. Application of this analysis to experimental data from the KcsA potassium channel identifies release as the RLS for outward permeation, consistent with available structural and computational evidence. Together, these results reveal how the physics of the electrochemical driving force shapes ion transport and provide an experimentally accessible strategy for connecting macroscopic electrophysiology measurements to microscopic transport mechanisms.

**Statement of significance:** One of the main challenges in the field of electrophysiology is connecting experimental currents from I-V data of ion channels to a detailed, atomistic mechanism of ion transport. We discovered that by examining the how the channel flux changes under different driving forces, we can identify the rate-, voltage-, and flux-limiting steps of ion transport. Understanding the relationships between ionic driving force and the corresponding channel behavior allows us to elucidate key details of the transport mechanism from macroscopic electrophysiology measurements.

## Introduction

Ion channels are essential regulators of ion homeostasis and electrical activity throughout the nervous, cardiovascular, and muscular systems. Dysfunction of these proteins has been linked to numerous human diseases, including cardiac arrhythmias, epilepsy, ataxia, diabetes, and chronic pain disorders (1,2). Due to their central physiological role, ion channels are important therapeutic targets for the treatment of neurological and cardiovascular disease. Ion channels are also commonly involved in off-target drug interactions, particularly hERG (3,4), leading to toxicity issues that can block the development of otherwise effective therapeutics (5). Understanding the molecular mechanisms governing ion selectivity, conduction, and gating is therefore critical for the development of more selective and effective pharmacological therapies.

Ion transport through many channels arises from the coordinated movement of ions through a narrow selectivity filter, where electrostatic interactions, hydration dynamics, and protein conformational fluctuations collectively determine conduction and selectivity (6–9). Although structural experiments, such as X-ray crystallography, and molecular dynamics (MD) simulations give remarkable insight into the molecular basis of ion channel function, electrophysiology is a primary experimental method for characterizing channel behavior under physiological conditions. By directly measuring ionic currents across membranes, electrophysiological techniques quantify key functional properties, including conductance, ion selectivity, gating, and voltage dependence. These measurements also reveal how changes in membrane voltage, environmental ion concentrations, and protein structure via disease-associated mutations alter channel properties (2,10), providing a critical link between molecular structure, physiological activity, and human disease.

While ion permeation describes the movement of ions through an open channel, channel gating and inactivation determine the probability that the channel occupies a conductive state. Depending upon the channel family, gating may be driven by membrane voltage (11), ligand binding (12), mechanical forces (13), temperature (14), or pH (7,15). Once open, the presence of ions within the channel pore or vestibule can stabilize the open conformation (16–18). As a result, ion permeation and inactivation can be coupled, with increased ion occupancy leading to longer dwell times in the open state (19,20). Coupling between permeation and inactivation is most evident in potassium channels, where ion occupancy within the selectivity filter influences the stability of the conductive state and the onset of C-type inactivation (21,22).

Among the diverse families of ion channels, potassium channels have emerged as one of the most extensively studied systems because they combine a relatively simple architecture with exceptional ion selectivity and rapid conduction. Despite sharing a highly conserved selectivity filter, members of the K^+^ channel family exhibit a wide range of conductances (23). Among these, a bacterial potassium channel, KcsA, is an important model for investigating ion permeation (18), selectivity (23,24), gating (17,25), and inactivation dynamics (16,21). As the first crystal structure of an ion channel (20), KcsA has been extensively characterized through structural biology, electrophysiology, and MD simulations. The abundance of structural and electrophysiological data makes KcsA an ideal system for directly relating experimentally measured currents to atomistic models of ion transport.

In our previous work (26) we demonstrated that current-voltage electrophysiology experiments (I-V curves) contain substantial information about the underlying permeation mechanism. Specifically, the locations of ion-binding sites and transition states—together with the relative rates of ion uptake, transfer, and release—produce characteristic current and conductance profiles (Fig. 1). These fingerprints enable researchers to identify the flux-limiting step (FLS) in the dominant mechanistic pathway and distinguish the physical origin of rectification with I-V and current-chemical potential (I-μ) curves. The FLS reflects the combined influence of the voltage-limiting step (VLS, defined as the transition with the lowest voltage-sensitivity), the rate-limiting step (RLS), the bulk ion concentration, and the applied membrane potential. While the VLS can be determined with MD simulations, as detailed in previous studies (27–29), identifying the RLS remains challenging.

**Figure 1.**
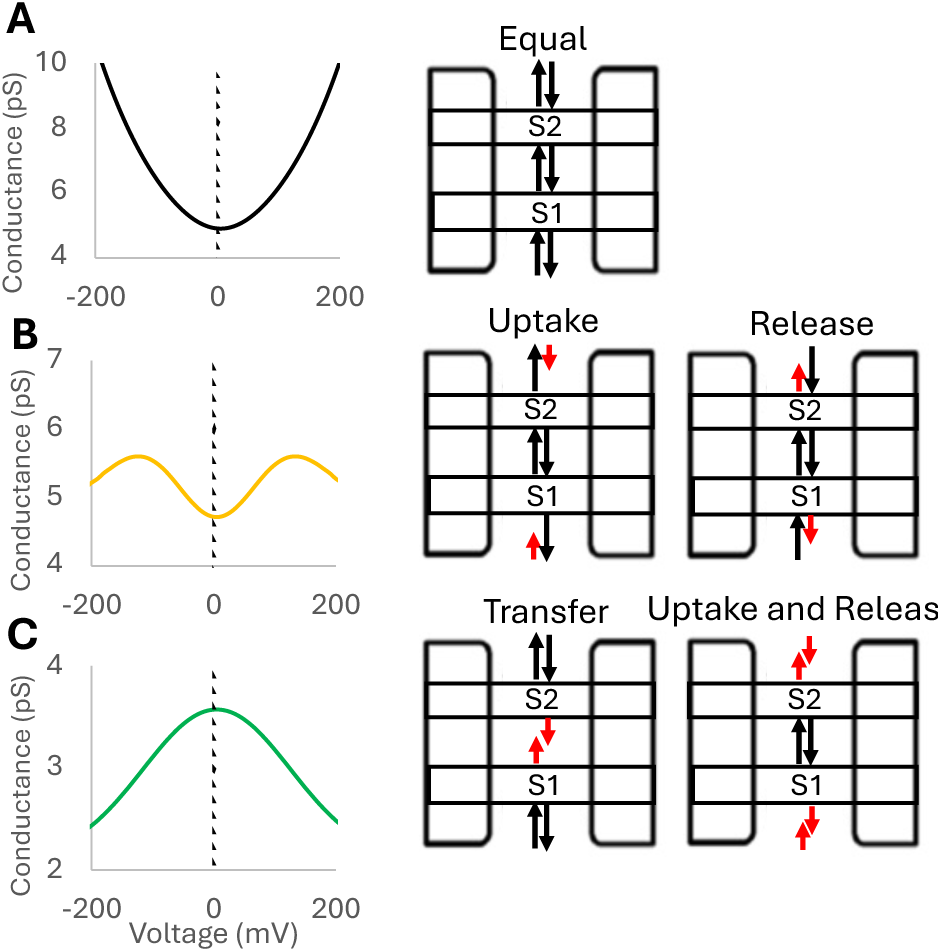
The conductance shape associated with (A) no FLS: shape 1, (B) uptake or release FLS: shape 2, or (C) transfer or equally uptake and release FLS: shape 3.

Our previous work also revealed that I-μ conductance changes shape with increasing chemical potential in uptake rate-limited model systems; however, release and transfer rate-limited systems remain indistinguishable (26). Unfortunately, I-μ curves are difficult to obtain experimentally, requiring many electrophysiological measurements controlled for net concentration at different chemical potential gradients. Furthermore, the VLS and RLS can oppose one another, producing a FLS that changes depending upon the magnitude of the voltage or chemical potential applied. This may be relevant in understanding the physiological role of many voltage-gated potassium channels, which function under strong outward chemical potential gradients that push K^+^ out of the cell against a negative voltage. Therefore, it would be ideal to identify a methodology capable of identifying the RLS across the range of electrochemical conditions under which a channel remains open, removing the confounding factor of voltage sensitivity, to clearly relate electrophysiological behavior to the underlying mechanism.

Here, we demonstrate how distinct effects of voltage and chemical potential gradients on ion permeation encode the RLS within the permeation pathway. By extending our previous work on I-V and I-μ analyses, we show that different flux profiles are produced when the RLS is uptake, transfer, or release due to how these transitions uniquely respond to changes in voltage and chemical potential. From these observations, we develop a practical electrophysiology protocol based on a small set of I-V relationships that clearly identify the RLS. Finally, we apply this methodology to KcsA to demonstrate its use and to provide new mechanistic insight into the RLS that governs outward ion conduction.

## Methods

### Electrochemically responsive kinetics

Ensemble averaged rates for rare-event transitions are determined by the probability of a system reaching the transition-state ensemble (sometimes described as a dividing surface) from a metastable reactant state, the frequency with which system dynamics approach the associated dividing surface, and the probability that crossing trajectories commit to product rather than recrossing to reactant. Within transition-state theory, this is commonly expressed as a dynamical prefactor (attempt frequency) multiplied by an exponential function of the free-energy, with an additional transmission coefficient accounting for recrossing dynamics near the transition-state ensemble. When considering the movement of a charged species through an open ion channel, the influence of electrochemical gradients should be considered for each of these terms. The presence of a transmembrane potential (V) influences all points along the transport free-energy surface,^27,30^ resulting in a new voltage-dependent free energy surface. In the kinetic modeling of open channel conduction, this new voltage-dependent free energy surface (G(z,V)) can be calculated using the descending fraction of the membrane potential (*φ*) that acts on a substrate as it traverses along the permeation pathway via the relationship (29):

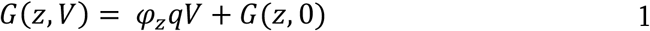

where q is the charge of the substrate, z is the position of the ion along the transport reaction coordinate, and *φ_z_* is the fraction of the membrane potential traversed between the start of the reaction coordinate and z. *φ_z_* can be computed using a number of different approaches (27,30), including continuum electrostatic potentials obtained from simulations under equal-magnitude positive and negative voltages as detailed in previous work (26,28,29,31,32).

For a transition from state i to state j, the voltage-modified Gibbs’ free energy of activation 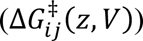 is given by the relationship:

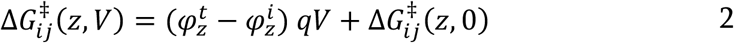

where 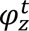 is the position of the transition state between states i and j and 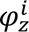 is the position of reactant basin of state i. Substitution of Eq. 2 into the Eyring-Polanyi equation (33), given by Eq. 3, produces the electrochemically responsive rate constants in Eq. 4 for each transition as follows:

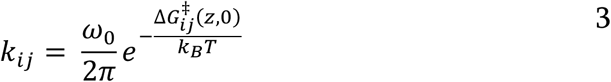

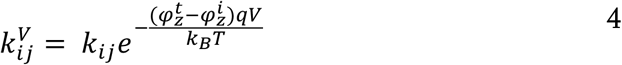

where *ω*_0_ is the attempt frequency, *k_B_* is Boltzmann’s constant, T is the temperature, and *k_i_*_j_ and 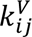 are the unmodified and voltage-modified rate constants, respectively. To account for the chemical potential driving ion translocation, the rates for ion uptake transitions are additionally altered by the bulk ionic activity, herein expressed as concentration ([*ion*]*_bulk_*), using a pseudo-first order approximation:

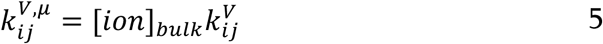

The flux (*J*) through this transition as a function of voltage can then be given by the relationship:

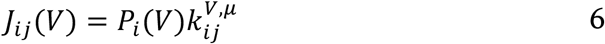

where *J_ij_* is the flux from state i to state j and *P_i_* is the probability of being in state i. Note that the attempt frequency and recrossing transmission coefficient can be influenced by voltage in a system-dependent manner and will be the focus of future work. The full derivation of electrochemically responsive rates is provided in previous work and recapitulated in the supporting material, Eq. S1-S14.

### MsRKM model construction

In previous work (26), we used model systems with two, three, or four stable binding sites and one of three symmetries: perfectly symmetric, in which equally spaced binding sites have identical transition rates and voltage sensitivities (Δ*φ* values); mirror symmetric, in which uptake/release/transfer parameters can vary but do so equivalently for inward and outward flux; or asymmetric, in which transition rates and their voltage sensitivities are allowed to be unequal for inward and outward flux, inducing rectification. In this study, we use the same labels for our kinetic model systems. Again, all model systems assume a linear voltage drop across the membrane (Fig. S1) and transition state locations are equidistant between stable binding sites unless noted otherwise in the text. For simplicity, the results shown herein use a two site model system; however, each trend was confirmed with three site and four site model systems, asymmetric transition state locations, and asymmetric voltage sensitivities to ensure our findings are independent of these facets. Specifications for all model systems used in this paper are provided (Tables S1-S2).

### I-V and I-μ analyses

As identified in our previous work (34), although both voltage and chemical potential driving forces can induce the same total permeation free-energy change, as quantified by the Nernst equation (35), voltage and chemical potential have different impacts on the underlying free energy surface of ion transport (Fig. 2). Chemical potential gradients only influence the pseudo first-order rates of uptake found in Eq. 5, while all non-uptake transitions maintain their zero-voltage rate. It is important to note that the physical origin of the effective increase in the free energy surface on the side with higher concentration is the increase in frequency that ions attempt to enter the channel. This falls out of the integration of the partition function of the reactant state (ions on one side of the membrane)— specifically the analytically differentiable translational freedom each ion has, or its free volume inversely proportional to concentration. This could in principle be represented by well widths on either side of the membrane, but the relative comparison between voltage and chemical potential is lost in this representation. Thus, we use the effective shift in free energy to capture the increase/decrease in ion uptake rates due to a chemical potential. The disproportionately larger impact on the side that decreases in concentration (Fig. 2 *A*) is due to the logarithmic relationship between chemical potential and free energy, as further explained below. In contrast, voltage influences every transition along the permeation pathway, with each binding site and transition state experiencing a distinct fraction of the transmembrane potential drop given by Eq. 4. This generally results in voltage and chemical potential gradients inducing distinct flux distributions through competing mechanistic pathways (28). We have demonstrated that conductance (*∂I*⁄*∂V*) shapes predict the FLS (*P_i_k_i_*_j_) for open channel conductance in a system-specific manner (26). In this paper, we extend that analysis to predict the RLS in the dominant flux pathway directly from I-V curves.

**Figure 2.**
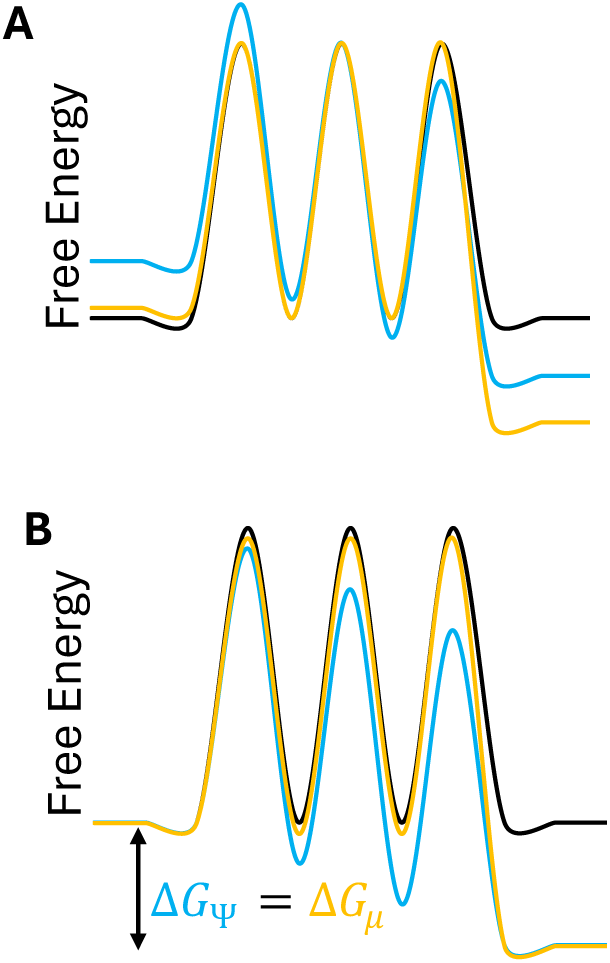
A free energy surface representing ion transport for a two site model channel (black). (A) Thermodynamic alignment highlights the unique effects of chemical potential (yellow) and voltage (blue) gradients. (B) Left alignment demonstrates that chemical potential and voltage produce equivalent net free energy differences.

## Results and discussion

### Distinct electrochemical sensitivities of ion uptake, transfer, and release

In previous studies, we observed that the flux driven by voltage was often larger than flux driven by Nernst-equivalent chemical potential gradients (26,28,29,34). However, further characterization of flux driven by chemical potential gradients (I-μ curves) reveals three different I-V and I-μ relationships that are correlated with the RLS (discussed below). To analyze these trends, we start by considering the different impacts of voltage and chemical potential gradients on specific ion transfer steps—broadly described by uptake, transfer, and release—in a perfectly symmetric model system (Fig. 3 *A*). These trends are demonstrated for outward flux, but hold for flux in either direction, noting that each direction is largely independent of the rates and trends for flux in the opposite direction (26).

**Figure 3.**
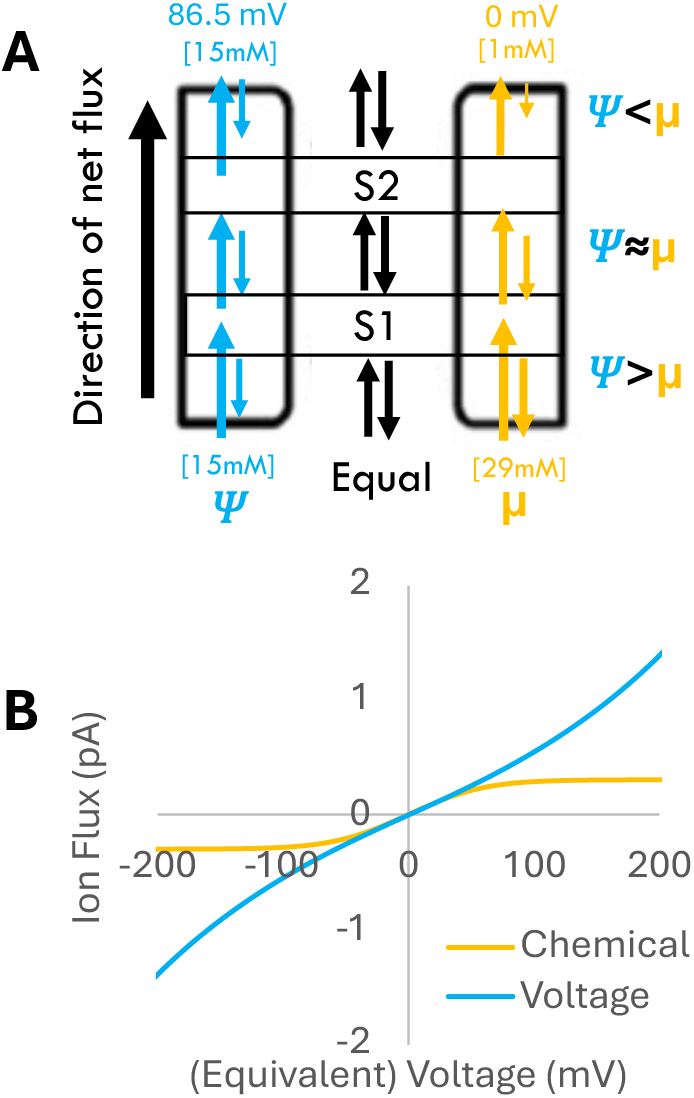
(A) The influence of voltage (blue) and chemical potential (yellow) on the bidirectional flux moving through a perfectly symmetric two site model system. (B) The difference in flux between an I-V and equivalent I-µ in a perfectly symmetric two site model system.

#### Uptake

Both voltage and chemical potential gradients increase the rate of uptake at S1, depicted at the bottom of the schematic in Fig. 3 *A*. However, voltage also slows the reverse step of ion release from S1, while the chemical potential gradient does not. As a result, voltage has a larger impact on uptake than the Nernst-equivalent chemical potential gradient.

#### Release

Voltage again increases the rate of release from site S2 (Fig. 3 *A*) while simultaneously decreasing the rate of reverse uptake. Again, the chemical driving force only slows reverse uptake due to low external ion concentration, but this decrease is proportionally larger than the increase in uptake to S1 (Fig. 2 *A*) due to the logarithmic expression in the Gibbs’ free energy of ion transport under an electrochemical potential:

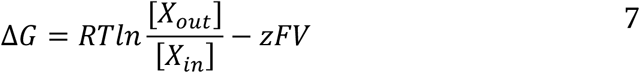

where [X_out_] and [X_in_] are the external and internal ion concentrations, respectively. Although the chemical potential gradient, which is given by the logarithmic term in Eq. 7, depends on the ratio of ion concentrations across the membrane, the proportion of the chemical potential change is not equivalent for uptake and release. The removal of ions from one side of the membrane has a disproportionately larger impact on Δ*G* than the addition of ions on the opposite side of the membrane due to the response of a logarithm to a fractional change under the constraint of mass conservation (the total number of ions must remain the same during the dissipation process). For example, consider moving 100 mM from a starting symmetric concentration of 300:300 mM ([*X_in_*]: [*X_out_*]) to create a 2/3 to 1/3 split of 400:200 mM. In this case, the chemical potential on the side that loses ions ([*X_out_*]) decreases by 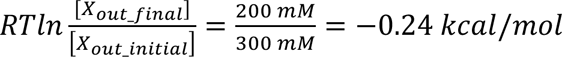 while the chemical potential on the side that gains ions ([*X_in_*]) increases by 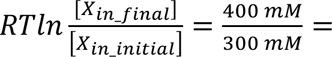 0.17 *kcal*/*mol*. Moreover, this difference becomes proportionally larger as the gradient increases (i.e., moving an additional 100 mM to create a 500:100 mM gradient yields an additional −0.41 kcal/mol outside and only +0.13 kcal/mol inside). Thus, the relative impact of moving ions across the membrane on Δ*G_μ_* is increasingly larger for the side that loses ions as dissipation continues. This results in chemical potential gradients having a larger influence on release, while voltage has a larger influence on uptake.

#### Transfer

Finally, the transfer flux between S1 and S2 near equilibrium was found to be relatively equal between the two driving forces, with voltage driving increased forward flux via a rate constant imbalance and chemical potential driving a similar increased forward flux via a population imbalance (Fig. 3 *A*). Collectively we find that ion uptake in the direction of net flux is more influenced by voltage, release is more influenced by chemical potential gradients, and transfer is approximately balanced in its response to voltage and chemical potential.

### Comparing I-V vs. I-μ analyses reveal the RLS

#### Uptake-limited model systems

Due to the above differences in electrochemical responses of uptake, transfer, and release, we find clear trends for I-V and I-μ curves that are indicative of the RLS. When uptake is rate limiting, (Fig. 4 *A*) we find that voltage-driven flux (Fig. 4 blue) exceeds flux produced by equivalent chemical potential gradient, even at low voltages (Fig. 4 yellow). This is consistent with uptake being more voltage sensitive. In addition to voltage speeding up the rate-limiting uptake to S1 while slowing reverse release, it also speeds transfer from S1 to S2 while slowing reverse transfer. This results in decreased occupancy at S1, which reduces blockage for faster forward flux. In contrast, while the rate-limiting uptake to S1 is sped up by a chemical potential gradient, there is no slowdown of back release and no change in the S1 to S2 transfer rate, leading to increased occupancy at S1, contributing to blockage and reverse flux (release from S1 to the inside). As discussed above, the relative impact of the increased internal concentration is further muted by the logarithmic nature of the chemical potential (Fig. 4 *C*). Collectively, when uptake is the RLS, voltage consistently drives greater flux than chemical potential gradients.

**Figure 4.**
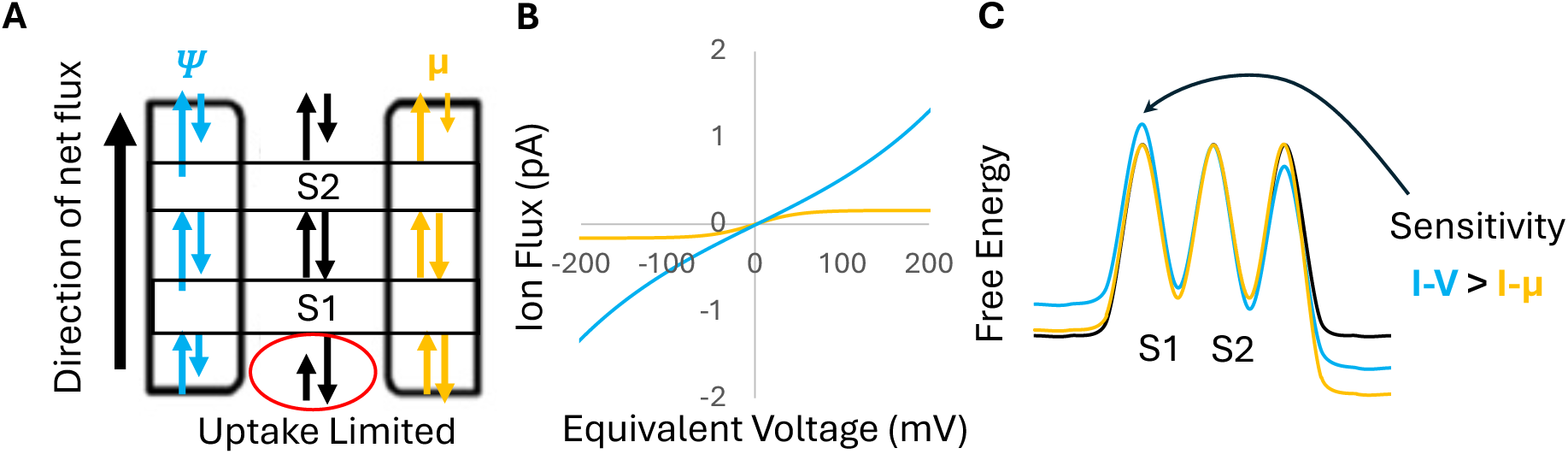
The influence of voltage (blue) and chemical potential (yellow) gradients on a two site model system with an uptake RLS. (A) Schematic displaying rate-constant differences between the two driving forces. (B) I-V and I-μ curves corresponding to the uptake-limited system in A. (C) Relative free energy differences between the driving forces.

#### Release-limited model systems

When release is rate limiting (Fig. 5), we find the opposite trend—greater flux is driven by the chemical potential gradient than by voltage. This is again consistent with release being more influenced by chemical potential, as discussed above. Voltage increases flux by speeding up release from S2 and decreasing reverse uptake. Voltage also increases forward flux from S1 to S2, which increases the population of S2 relative to the population from chemically driven flux, which further supports release flux. However, the impact of the chemical potential gradient on slowing down reverse uptake is more significant than the changes due to voltage. Thus, for release-limited model systems, chemical potential gradients drive larger flux than Nernst-equivalent voltage up to a given cross-over point when the former plateaus due to the logarithmic nature of the entropic chemical driving force (at ∼±140 mV in Fig. 5 *B* and further discussed below) and voltage once again drives more flux as it continues to change rates throughout the energy surface.

**Figure 5.**
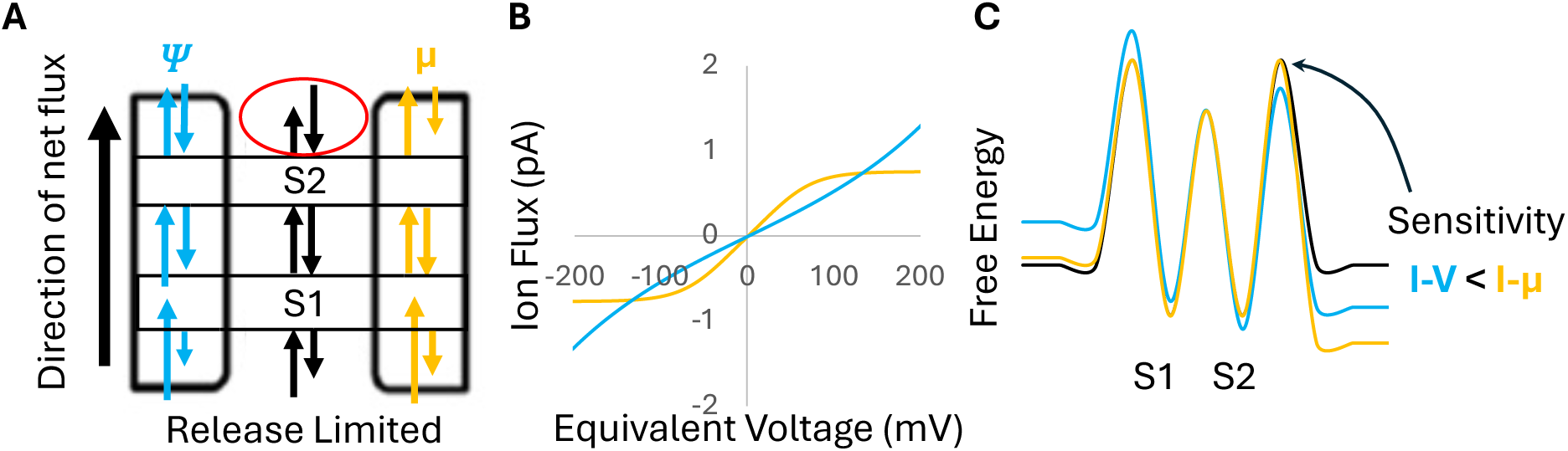
The influence of voltage (blue) and chemical potential (yellow) gradients on a two site model system with a release RLS. (A) Schematic displaying rate constant differences by arrow length for the two driving forces. (B) I-V and I-μ curves corresponding to the release-limited system in A. (C) Relative free energy differences between the driving forces.

#### Transfer-limited model systems

Finally, when transfer is rate limiting (Fig. 6) or when there is no dominant RLS (Fig. 3), there is a region where the observed flux is similar between the driving forces, again with the caveat that voltage-driven flux will eventually exceed chemical potential-driven flux. This is again consistent with the above findings, where the equivalence for small to modest driving forces comes from similar effects of voltage increasing forward flux via rate changes and chemical potential increasing forward flux via population changes. However, for sufficiently large gradients, voltage induces greater flux than the equivalent chemical potential gradient (Fig. 6 *B*).

**Figure 6.**
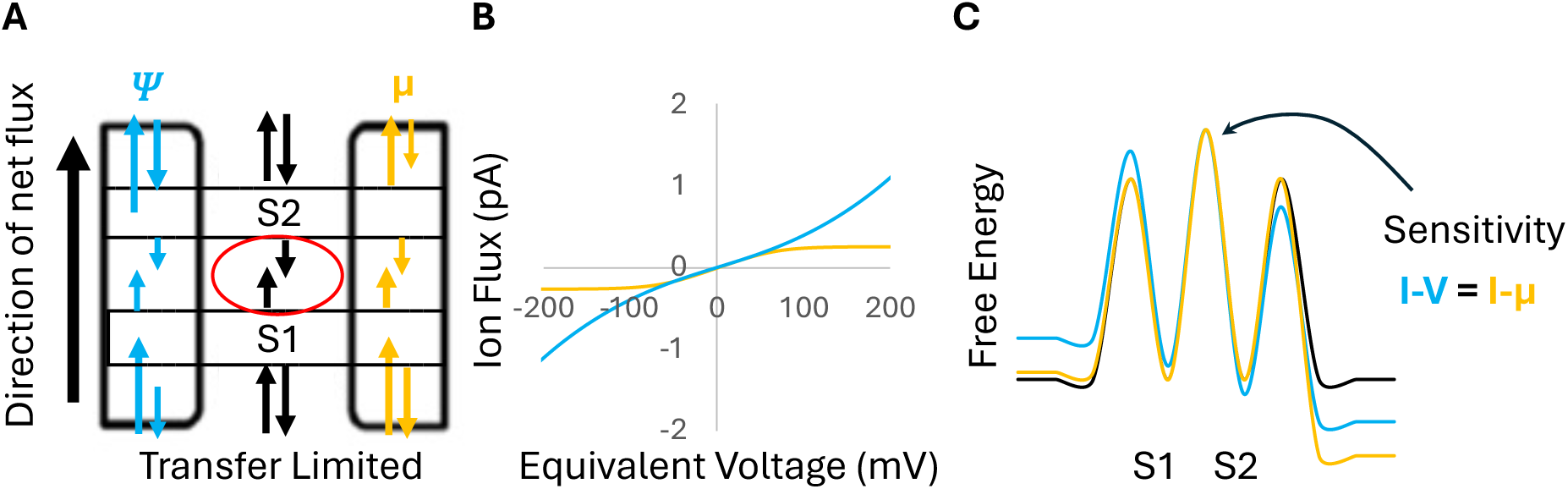
The influence of voltage (blue) and chemical potential (yellow) gradients on a two site model system with a transfer RLS. (A) Schematic displaying rate constant differences with arrow length for the two driving forces. (B) I-V and I-μ curves corresponding to the transfer-limited system in A. (C) Relative free energy differences between the driving forces.

#### Chemical potential-driven flux plateaus

As the driving force increases in magnitude, voltage-driven flux eventually overtakes the plateauing chemical potential-driven flux, even in release- and transfer-limited systems. This is because the ability of chemical potential to increase flux via population alteration is ultimately constrained by two factors: diminishing returns in concentration changes due to its logarithmic nature, and the inherent rate-limiting steps in the transport process. In other words, chemical potential-driven flux plateaus because the entropic driving force for transport is logarithmic, reflecting the concave nature of the entropy of mixing, wherein each additional mixing increment has a smaller impact on the side with larger concentrations. Additionally, biological transport systems add an additional layer of saturation because their microscopic transition rates are finite, causing the transport cycle itself to become rate limiting. As the chemical potential gradient increases, the population of the binding site preceding the RLS saturates, and flux becomes limited by a RLS that is independent of the chemical potential driving force. In contrast, voltage is unconstrained—modifying the free energy surface with an exponential relationship and continuing to speed up the RLS with increasing voltage regardless of its location in the transport cycle. Thus, voltage takes over when the flux driven by chemical potential plateaus. Consistent with this logic, the region of overlap between the I-V and I-μ curves depends on the degree to which transfer is rate limiting. Model systems with much slower transfer rates have larger regions of overlap (Fig. S2) and requires larger voltages to bring the relative rate of the RLS down enough for voltage to overtake chemical potential-driven flux.

### Extracting the RLS direction from I-V assays

#### Measured vs. predicted flux analysis

Although the RLS can be visually extracted by comparing I-V and I-µ curves, as described above (Figs. 4 *B*, 5 *B*, and 6 *B*), it is generally experimentally difficult to obtain I-µ curves. Additionally, these trends can be obfuscated when the voltage sensitivity counteracts the RLS. For example, Fig. 7 shows a model system in which release is the RLS and also the most voltage responsive, with increasing voltage speeding up release. The origin of this sensitivity is a transition state that is closer to extracellular solution than the last binding site, giving extracellular release a larger Δ*φ* than extracellular uptake (Table S2). This combination obscures the RLS in the +mV flux direction, as it produces a region of overlap between the I-V and I-μ curves, characteristic of a transfer-limited process (Fig. 7 *A*).

**Figure 7.**
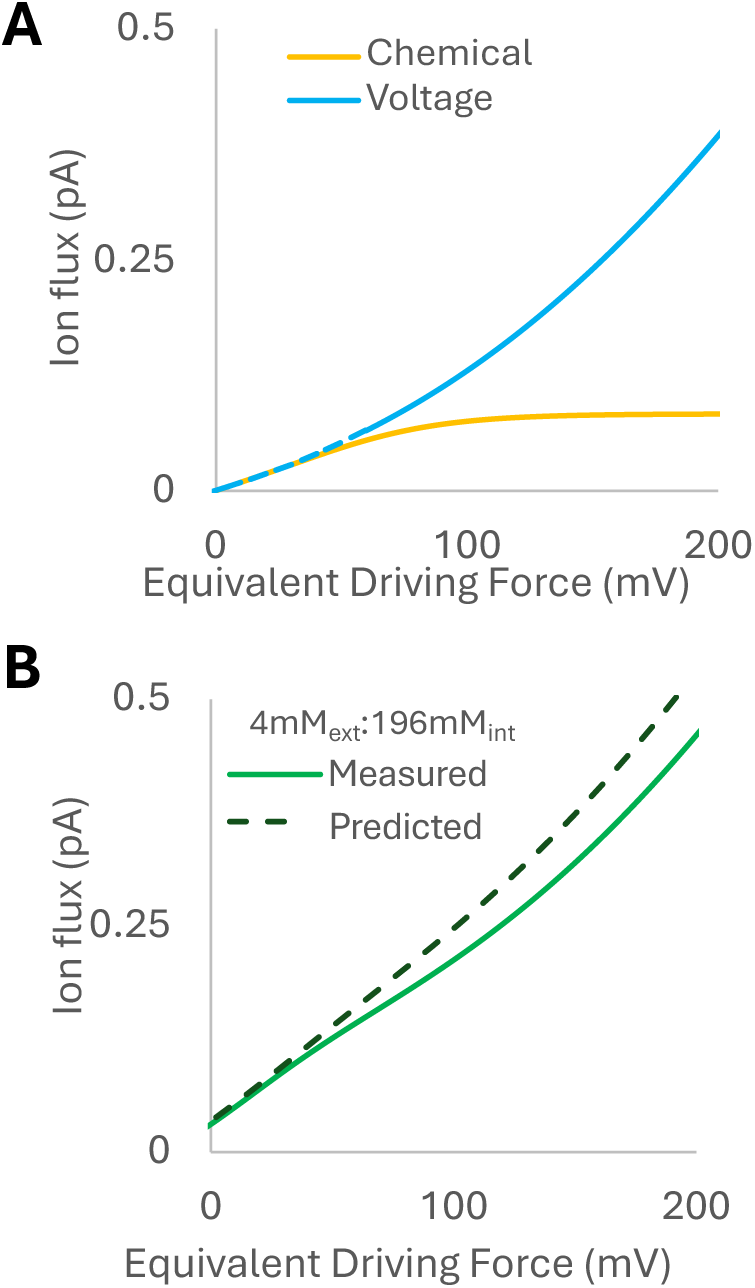
(A) I-V (blue) and I-μ (yellow) curves show a transfer RLS instead of the correct release RLS. (B) Comparing Measured (black dashed) and Predicted (green solid) I-V curves reveal the correct RLS.

Fortunately, the RLS of the dominant flux producing pathway can be cleanly extracted from the combination of three standard I-V curves (Fig. 8). These I-V curves compare the flux from the simple summation of the pure driving forces, which we call ‘Predicted,’ to the flux actually driven when the two forces act together, which we call ‘Measured.’ The first I-V assay should be performed with symmetric concentrations ([X_in_] = [X_out_]). The second should be performed with a constant outward gradient ([X_in_] > [X_out_]) to characterize outward flux, while the third should have an inward gradient ([X_in_] < [X_out_]) to characterize inward flux. Ideally, each assay should have the same total concentration ([X_tot_] = [X_in_] + [X_out_]), although the essential trends seem to be tolerant of modest differences (up to ∼10%) in net concentration (Fig. S3).

**Figure 8.**
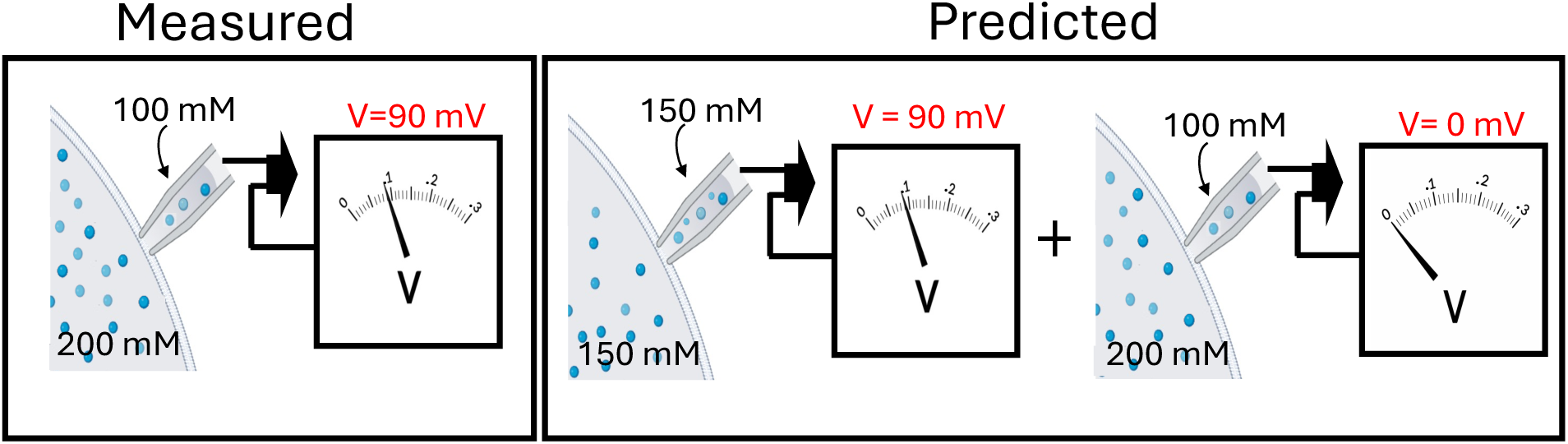
The two experiments required to produce Measured and Predicted I-V curves for outward K^+^ flux. Measured: Experimental I-V curve with a constant outward chemical potential gradient (200:100 mM [K^+^_int_]: [K^+^_ext_]). Predicted: A symmetric concentration I-V curve plus the flux at 0 mV of the Measured I-V curve.

Below we describe ‘Measured vs. Predicted’ analysis for outward flux (+mV), noting the same trends and logic hold for inward flux. The Predicted flux is calculated by adding the flux values obtained at each voltage from the symmetric I-V curve to the flux obtained at 0 V with the asymmetric gradient (Fig. 8).

#### Uptake RLS

When uptake is the RLS, the Measured I-V curve shows increasingly more flux than the Predicted I-V curve (Fig. 9 *A*). This is a result of the combined driving forces working synergistically to accelerate the rate-limiting uptake, while voltage is still suppressing reverse release. When voltage and chemical potential are applied together (Measured), the chemical potential gradient increases the population of the ‘reactant’ (ions inside) attempting to cross the RLS. This leads to increased flux relative to the summation of flux driven by voltage and a chemical potential gradient independently.

**Figure 9.**
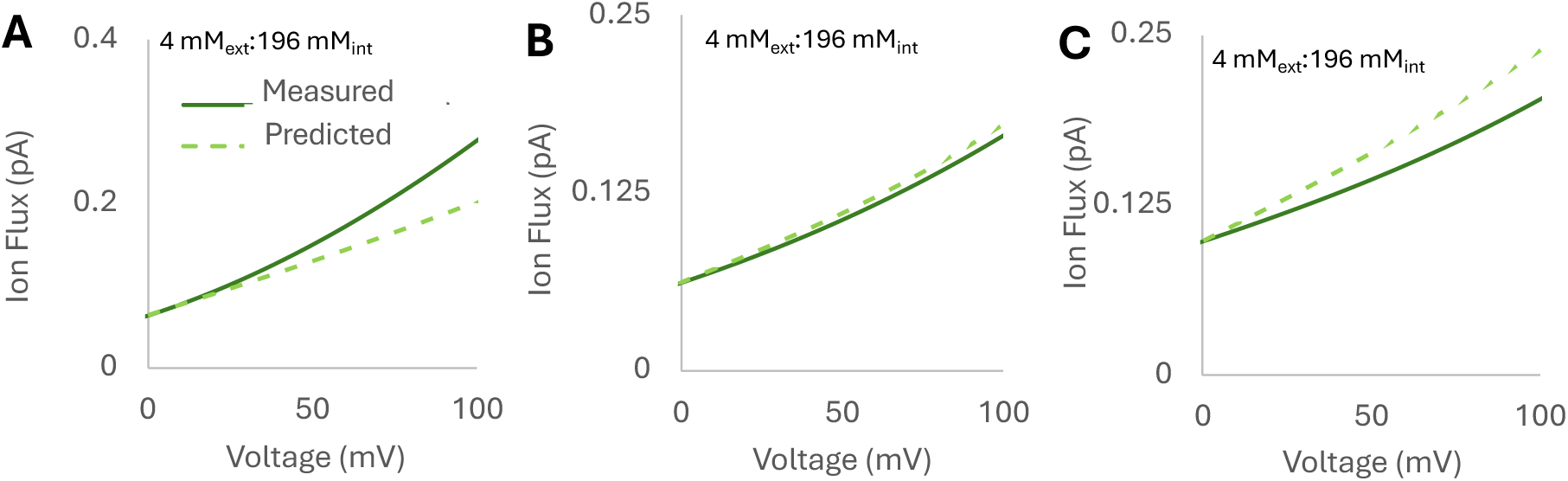
Measured (solid) vs. Predicted (dashed) flux for (A) uptake-limited, (B) transfer-limited, and (C) release-limited two site model systems.

#### Release RLS

The opposite is observed for model systems in which release is the RLS (Fig. 9 *C*). Here the two forces act in opposition, with the suppression of reverse uptake at S2 by the low outside concentration decreasing the population of the ‘reactant’ (ions at S2) before the RLS, resulting in lower actual (Measured) flux compared to a summation of the fluxes driven independently (Predicted).

#### Transfer RLS

Finally, we find that transfer-limited systems (Fig. 9 *B*) show similar Measured and Predicted I-V curves, consistent with the equivalent effects of voltage and chemical potential on transfer-limited systems described above. Collectively, we find the actual experimental flux (Measured) will be larger than that anticipated from their summation (Predicted) for uptake-limited processes, less for release-limited processes, and equivalent for transfer-limited processes

### Application of flux and Measured vs. Predicted analyses to KcsA

We demonstrate our methodological pipeline for the KcsA potassium channel using experimental asymmetric I-V curve from Zakharian et al. (36), (10 mM [K^+^_ext_] pH 7: 200 mM [K^+^_int_] pH 7), and experimental symmetric I-V curve from Nimigean et al. (37), (100 mM [K^+^_ext_] pH 6: 100 mM [K^+^_int_] pH 7). This analysis assumes that gating effects have been correctly removed from the I-V analysis in both experiments and that the difference in internal pH does not alter the channel’s open channel conductance, as further supported below.

#### Predictions for KcsA

Our methodological pipeline begins with analysis of the FLS for both inward and outward current, revealed by the conductance shape of the symmetric I-V curve (Fig. 10 *B*), followed by analysis of the RLS for outward flux.

**Figure 10.**
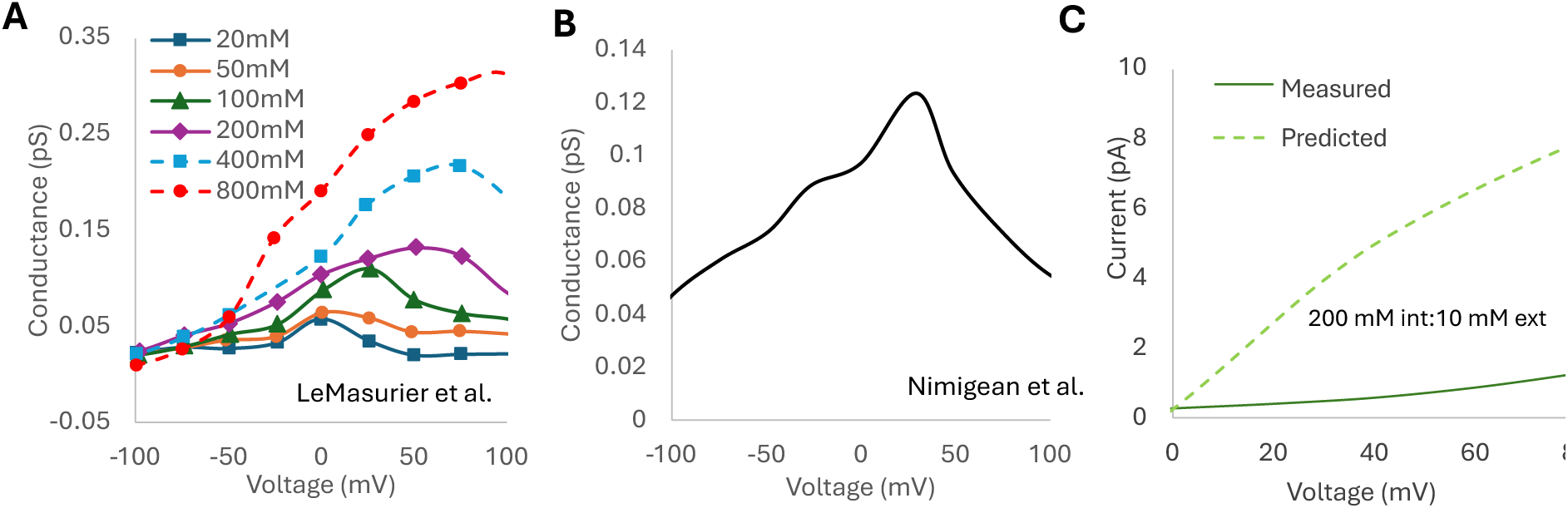
Extracting mechanistic insight for KcsA using experimental I-V curves. (A) Conductance shape evolution across a range of symmetric concentrations. (B) The symmetric I-V curve used for Measured vs. Predicted analysis shows conductance shape 3 for inward flux and conductance shape 2 for outward flux. (C) Comparing the Measured (solid) vs. Predicted (dashed) flux reveals the former is smaller—indicating release is the RLS for outward flux.

#### Inward flux (-mV)

For inward flux, we see conductance shape 3 from the symmetric I-V curve (Fig. 10 *B*), indicating that the FLS is either transfer or equally uptake and release. Additional I-V curves from symmetric 20 mM to 800 mM concentrations all retain shape 3 (Fig. 10 *A*), which indicates that flux is most likely limited by transfer at 100 mM [K^+^_ext_] since there is no change in shape for increasing or decreasing concentration, although the range of concentrations (20-800 mM) does not exclude the possibility of binding and release together being flux limiting.

#### Outward flux (+mV)

Turning to outward flux, the symmetric I-V curve (Fig. 10 *B*) fits conductance shape 2, suggesting either uptake or release are flux limiting. However, the location of the maximum in outward flux shifts from approximately +30 mV when [K^+^_int_] is 100 mM (Fig. 10 *B*) to +50 mV when [K^+^_int_] is 200 mM (Fig. 10 *A*). For uptake or release rate-limited systems, the location of the conductance shape 2 peak reflects the disparity between the uptake and release fluxes, quantified by their ratio. As uptake and release fluxes become more nearly equal, the profile transitions to a different conductance shape. Because increasing [K^+^_int_] from 100 mM to 200 mM shifts the peak farther from 0 mV—indicating a larger imbalance—the flux must be release limited at both concentrations. As [K^+^_int_] is reduced and the uptake flux correspondingly decreases, the conductance peak moves further in, transitioning at 50 mM to conductance shape 3. For 100 mM and above, outward flux is clearly release-limited.

To determine the RLS, we obtain the Predicted flux by summing the pure voltage-driven flux in the symmetric I-V curve (37) with the pure chemical potential-driven flux at 0 mV in the asymmetric I-V curve (36) (Fig. 10 *C*). The Measured flux is simply the asymmetric I-V curve. Importantly, these two sets of experimental data have similar net concentrations (210 mM for Zakharian et al. (36) and 200 mM for Nimigean et al. (37)), which is essential for the Measured vs. Predicted analysis. However, since the analysis is direction dependent, we can only assess the RLS for outflux.

We find that the Predicted flux is substantially larger than the Measured flux, indicating release is the RLS. Although the RLS and FLS don’t have to be the same step, they seem to both be release for outward flux in KcsA at 100 mM [K^+^_int_]. Based on the analyses above, a release RLS indicates that KcsA in the open conformation is more sensitive to changes in chemical potential than to equivalent voltage changes (Fig. 5). This prediction is interesting in the context of the well-known dependence of C-type inactivation in KcsA on external K^+^ concentrations.

### Comparing to previous work on KcsA

#### Computational experiments on wild type KcsA

Although MD simulations are subject to force field errors and other approximations that generally produce K^+^ channels’ currents that are inconsistent with electrophysiology experiments, simulations still play an invaluable role in unravelling the molecular mechanisms of ion channel gating and permeation. Simulations of Allen et al. (38), demonstrated that multiple ion occupancies are required to release potassium from the selectivity filter. This supports the claim that KcsA flux is release-limited as the energetic barrier of release is high enough to be inaccessible without the additional energetic push from the electrostatic repulsion between additional K^+^ ions (39–41). One of the biggest challenges in simulating wild type KcsA is that the selectivity filter can collapse into a nonconductive conformation (C-type inactivation). To overcome this complication, many computational and electrophysiological studies instead employ the E71A mutant, which suppresses C-type inactivation. We observe conductance shape 3 from analysis of experimental I-V data for E71A at 100 mM for the +mV direction (Fig. S5 *A*), which indicates a FLS of either transfer or equally uptake and release flux limited. Conductance analysis of the recent results of ECC-corrected simulations (42) also shows conductance shape 3 (at 100 mM) with a shift to conductance shape 2 (release-limited) at ∼500 mM (Fig. S5 *A*). This suggests that E71A responds similarly to concentration as wild type KcsA but reaches nearly equivalent uptake and release fluxes at higher concentrations (∼100 mM vs. 20 mM). Similarly, recent Markov modeling of KcsA E71A mutant by Furini et al. (43) also appears to report release-limited rate constants at 0 mV.

#### Experimental results of KcsA

Crystal structures of wild-type KcsA have a notable shift in ion occupancy between 150 mM symmetric K^+^ (1K4C) and 3 mM symmetric K^+^ (1K4D) (44). At 150 mM, density was interpreted as two ion occupancies within the selectivity filter with equal proportionality. This proportional split of ion occupancy was corroborated later by ssNMR (45) and 2D-IR (46, 47) data. Since these observed ion occupancies are split between the top and bottom section of the selectivity filter, this establishes the two regions have similar stability under zero voltage conditions. More detailed ssNMR of wild type KcsA at different concentrations and with a fully open activation gate at pH 3.0 showed a notable increase in the probability of occupancy of Scav, S3, and S4 sites once the symmetric K^+^ concentration was increased to, and above, 80 mM (48). We infer that these occupancy trends at the bottom of the selectivity filter are similar to trends at the top of the selectivity filter, which was not observable in this latter study. This occupancy change is consistent with the conductance shift to conductance shape 2 (likely release-limited) observed in Fig. 10 *A* between 50 mM and 100 mM. At concentrations lower than 100 mM, the rate of binding slows and competes with release to destabilize an ion occupied selectivity filter, as seen in previous studies (45, 48–51). At a pH of 7, this shift notably occurs at lower concentrations (10 μM), consistent with allosteric destabilization of the selectivity filter by pH-triggered gating. However, the measurements of Nimigean et al. (37) show that KcsA retains both conductance shape 2 and its peak position for outward flux over the pH range 3.0–6.0 for symmetric 100 mM conditions (Fig. S4 *A*), suggesting that pH does not influence the balance between uptake and release in open channel conductance. Additionally, the ss-NMR data of the H25R/E118A double mutant, which maintains an open pH gate across the same range of pH values, shows the same increased occupancy of Scav, S3, and S4 at 80 mM as wild type KcsA at a pH 3.0 (50). This suggests that while pH-related gating shifts the relative stability of the selectivity filter and channel occupancies, it does not change the flux-limiting step of open channel outflux prior to inactivation. Above an internal concentration of ∼80 mM, release is flux limiting for outward flux.

### Conclusions

In this work, we extend our efforts to extract mechanistic information directly from electrophysiology measurements of current-voltage relationships. In our previous work, we showed that characteristic conductance shapes encode whether uptake, transfer, release, or a combination of these transitions carries the least flux (the FLS) during ion conduction. Here, we extend that framework by showing that a combination of three standard current-voltage measurements can identify the underlying kinetically slowest transition (the RLS). Together, these complementary analyses enable both the transitions carrying the least flux and the kinetically slowest transition to be identified directly from current-voltage measurements, providing a more complete mechanistic interpretation of ion-channel electrophysiology.

Critical to these analyses are the different physical mechanisms by which voltage and chemical potential perturb ion transport. Voltage reshapes the free-energy landscape throughout the permeation pathway, altering every transition rate, whereas chemical potential primarily changes the probability that ions attempt uptake into the channel. Consequently, uptake, transfer, and release respond differently to the two driving forces.

This difference leads to several important physical consequences. First, uptake-rate-limited transport is intrinsically more responsive to voltage because voltage both accelerates uptake and suppresses reverse release, whereas chemical potential gradients only increase the frequency of uptake attempts. In contrast, release-limited transport is more responsive to chemical potential because the logarithmic dependence of the entropy of mixing causes decreases in concentration on the depleted side of the membrane to have a disproportionately larger effect than equivalent increases on the enriched side. Thus, voltage drives larger flux than Nernst-equivalent chemical potential gradients for uptake-limited systems, whereas chemical potential gradients drive larger flux for release-limited systems. Transfer-limited systems respond similarly to both driving forces.

A second important consequence is that chemical potential-driven flux is fundamentally bounded, whereas voltage-driven flux is not. As chemical potential gradients increase, the logarithmic chemical potential produces progressively smaller increases in driving force while the population preceding the RLS approaches saturation. Once this occurs, additional increases in concentration produce diminishing increases in current because transport becomes limited by the intrinsic kinetics of the rate-limiting transition. Voltage, however, continues to modify transition barriers throughout the permeation pathway and therefore continues to accelerate transport as the applied potential increases. This explains why flux driven by a chemical potential plateaus at high potentials, while that driven by voltage continues to increase for all three classes of transport mechanisms.

Finally, these physical principles provide a practical method for identifying the RLS from conventional electrophysiology experiments. Because direct I-μ measurements are experimentally challenging and voltage asymmetry can obscure the underlying RLS, we developed the Predicted vs. Measured analysis using only three standard I-V experiments. When uptake is rate limiting, voltage and chemical potential act synergistically, producing Measured currents larger than those Predicted from the sum of their independent effects. When release is rate limiting, the two perturbations partially oppose one another, producing Measured currents smaller than those Predicted. Transfer-limited systems exhibit little difference between the two. Application of this framework to published KcsA data predicts that open channel outward currents are both flux and rate limited by release at higher concentrations but switch to being both binding and release limited at lower concentrations, consistent with previous computational and structural studies. We additionally predict inward current is transfer flux limited, but additional assays are needed to determine the RLS for inward currents.

Collectively, these findings demonstrate that differences between voltage- and chemical potential-driven fluxes encode mechanistic information about the transport process itself. By combining conductance-shape analysis with the Predicted vs. Measured framework, routine electrophysiological measurements can be used to identify both the FLS and RLS under specific electrochemical conditions, providing a direct bridge between macroscopic transport measurements and microscopic transport mechanisms.

## Author contributions

H.W.D., R.C., and J.M.J.S. designed the research. H.W.D. and O.H. built all kinetic models and H.W.D. validated and analyzed the data from all models used within this paper. H.W.D, O.H., and A.D.D. ran the kinetic model systems. All authors helped interpret the final results. J.M.J.S. directed and funded the research. All authors wrote the manuscript and have given approval on the final version.

## Acknowledgments

This work was supported by the NIH NIGMS (R35GM143117) and the computational resources provided by Expanse at the San Diego Supercomputing Center through the Advanced Cyberinfrastructure Coordination Ecosystem: Services and Support (ACCESS) program (allocation MCB200018) supported by NSF (Grant Nos. 2138259, 2138286, 2138307, 2137603, and 2138296), as well as the Center for High-Performance Computing (CHPC) at the University of Utah.

## Supporting material

The following files are available free of charge.

Supplementary material file includes additional derivations found in Eq. S1-S14, MsRKM model specifications (Tables S1-S2), and figures (Figs. S1-S5). An example voltage coupling profile can be found in Fig. S1. Results include additional I-V vs. I-μ comparison between transfer limited models (Fig. S2), an assessment of the accuracy of the Predicted vs. Measured analysis at unequal net concentrations (Fig. S3), conductance shape analysis of KcsA at different symmetric pH (Fig. S4), and conductance shapes for wild type KcsA, the E71A mutant, and ECC corrected E71A of Hui et al. (42) (Fig. S5).

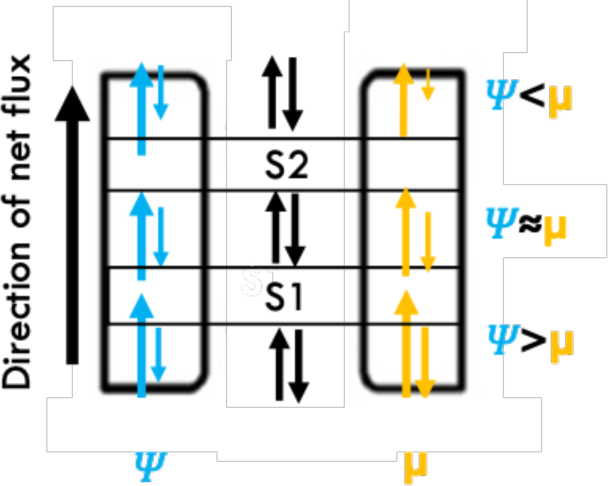
For Table of Contents only

## References

(1) Hryciw, D. H. 2024. Special Issue: “Recent Advances in Ion Channels and Ion Channelopathies”. Int. J. Mol. Sci. 25:3598.

(2) Harraz, O. F.; Delpire, E. 2024. Recent insights into channelopathies. Physiol. Rev. 104:23–31. DOI: 10.1152/physrev.00022.2023.

(3) Kalyaanamoorthy, S.; Barakat, K. H. 2018. Development of Safe Drugs: The hERG Challenge. Med. Res. Rev. 38:525–555. doi: 10.1002/med.21445.

(4) Farm, H. J.; Clerx, M.; Cooper, F.; Polonchuk, L.; Wang, K.; Gavaghan, D. J.; Lei, C. L. 2023. Importance of modelling hERG binding in predicting drug-induced action potential prolongations for drug safety assessment. Front. Pharmacol. 14. doi: 10.3389/fphar.2023.1110555.

(5) Lopez-Mateos, D.; Harris, B. J.; Hernández-González, A.; Yarov-Yarovoy, V.; Wulff, H. 2025. Recent advances in the pharmacology of voltage-gated ion channels. Pharmacol. Rev. 77:100090. doi: 10.1016/j.pharmr.2025.100090.

(6) Roux, B. 2017. Ion channels and ion selectivity. Essays Biochem. 61:201–209. doi: 10.1042/ebc20160074.

(7) Lee, J.; Kang, M.; Kim, S.; Chang, I. 2020. Structural and molecular insight into the pH-induced low-permeability of the voltage-gated potassium channel Kv1.2 through dewetting of the water cavity. PLoS Comput. Biol. 16:e1007405. doi: 10.1371/journal.pcbi.1007405.

(8) Ryan, M. J.; Gao, L.; Valiyaveetil, F. I.; Kananenka, A. A.; Zanni, M. T. 2024. Water inside the Selectivity Filter of a K(+) Ion Channel: Structural Heterogeneity, Picosecond Dynamics, and Hydrogen Bonding. J. Am. Chem. Soc. 146:1543–1553. doi: 10.1021/jacs.3c11513.

(9) Bosio, S.; Gazzoni, D.; Domene, C.; Masetti, M.; Furini, S. 2026. Role of Water Models in Simulations of Ion Conduction in Potassium Channels. J. Chem. Theory Comput. 22:1177–1186. doi: 10.1021/acs.jctc.5c01787.

(10) Mironenko, A.; de Groot, B. L.; Kopec, W. 2024. Selectivity filter mutations shift ion permeation mechanism in potassium channels. PNAS Nexus. 3. doi: 10.1093/pnasnexus/pgae272.

(11) Davila-Velderrain, J.; van Giesen, L. 2025. Voltage-gated ion channel diversity underlies neuronal excitability and nervous system evolution. Nat. Commun. 16:11534. doi: 10.1038/s41467-025-66641-8.

(12) Tovar, K. R.; Westbrook, G. L. 2026. Chapter 23 - Ligand-gated ion channels. In Cell Physiology Source Book (Fifth Edition), Alvarez-Leefmans, F. J., Delpire, E., Kaneshiro, E. S. Eds.; Academic Press, 557–572.

(13) Ranade, Sanjeev S.; Syeda, R.; Patapoutian, A. 2015. Mechanically Activated Ion Channels. Neuron. 87:1162–1179. doi: 10.1016/j.neuron.2015.08.032.

(14) Cesare, P.; Moriondo, A.; Vellani, V.; McNaughton, P. A. 1999. Ion channels gated by heat. Proc. Natl. Acad. Sci. U. S. A. 96:7658–7663. doi: 10.1073/pnas.96.14.7658.

(15) Thompson, A. N.; Posson, D. J.; Parsa, P. V.; Nimigean, C. M. 2008. Molecular mechanism of pH sensing in KcsA potassium channels. Proc. Natl. Acad. Sci. U. S. A. 105:6900–6905. doi:10.1073/pnas.0800873105.

(16) LeMasurier, M.; Heginbotham, L.; Miller, C. 2001. KcsA: it’s a potassium channel. J. Gen. Physiol. 118:303–314. doi: 10.1085/jgp.118.3.303.

(17) Baker, K. A.; Tzitzilonis, C.; Kwiatkowski, W.; Choe, S.; Riek, R. 2007. Conformational dynamics of the KcsA potassium channel governs gating properties. Nat. Struct. Mol. Biol. 14:1089–1095. doi: 10.1038/nsmb1311.

(18) Tilegenova, C.; Cortes, D. M.; Cuello, L. G. 2017. Hysteresis of KcsA potassium channel’s activation & deactivation gating is caused by structural changes at the channel’s selectivity filter. Proc. Natl. Acad. Sci. U. S. A. 114:3234–3239. doi:10.1073/pnas.1618101114.

(19) Baukrowitz, T.; Yellen, G. 1996. Use-Dependent Blockers and Exit Rate of the Last Ion from the Multi-Ion Pore of a K+ Channel. Science. 271:653–656. doi:10.1126/science.271.5249.653.

(20) Doyle, D. A.; Cabral, J. M.; Pfuetzner, R. A.; Kuo, A.; Gulbis, J. M.; Cohen, S. L.; Chait, B. T.; MacKinnon, R. 1998. The Structure of the Potassium Channel: Molecular Basis of K+ Conduction and Selectivity. Science. 280:69–77. doi:10.1126/science.280.5360.69.

(21) Xu, Y.; McDermott, A. E. 2019. Inactivation in the potassium channel KcsA. J. Struct. Biol: X. 3:100009. doi: 10.1016/j.yjsbx.2019.100009.

(22) Reddi, R.; Matulef, K.; Riederer, E. A.; Whorton, M. R.; Valiyaveetil, F. I. 2022. Structural basis for C-type inactivation in a Shaker family voltage-gated K+ channel. Sci. Adv. 8:eabm8804. doi:10.1126/sciadv.abm8804.

(23) Lee, S.-J.; Schillings, J.; De Vries, R.; Matamoros, M.; De Groot, B. L.; Nichols, C. G. 2025. From selective permeation to physiology in potassium channels. Function. 6:zqaf050.

(24) Thompson, A. N.; Kim, I.; Panosian, T. D.; Iverson, T. M.; Allen, T. W.; Nimigean, C. M. 2009. Mechanism of potassium-channel selectivity revealed by Na(+) and Li(+) binding sites within the KcsA pore. Nat. Struct. Mol. Biol. 16:1317–1324. doi: 10.1038/nsmb.1703.

(25) Catacuzzeno, L.; Conti, F.; Franciolini, F. 2023. Fifty years of gating currents and channel gating. J. Gen. Physiol. 155. doi: 10.1085/jgp.202313380.

(26) Weckel-Dahman, H.; Carlsen, R.; Daum, A.; Swanson, J. M. J. 2026. From Flux to Function: Extracting Mechanistic Insights from Ion Channels via I–V and I–μ Analyses. J. Phys. Chem. B. 130:2733–2742. doi: 10.1021/acs.jpcb.5c06019.

(27) Roux, B. 2008. The Membrane Potential and its Representation by a Constant Electric Field in Computer Simulations. Biophys. J. 95:4205–4216. doi: 10.1529/biophysj.108.136499.

(28) Weckel-Dahman, H.; Carlsen, R.; Daum, A.; He, M.; Southam, T. G.; Swanson, J. M. J. 2025. Ion Channel Reaction Networks: Dielectric Screening and the Importance of Off-Pathway Flux. J. Chem. Theory Comp. 21:3700–3711. doi: 10.1021/acs.jctc.4c01569.

(29) Weckel-Dahman, H.; Carlsen, R.; Swanson, J. M. J. 2024. Multiscale Responsive Kinetic Modeling: Quantifying Biomolecular Reaction Flux under Varying Electrochemical Conditions. J. Chem. Theory Comp. 21:374–389. doi: 10.1021/acs.jctc.4c00872.

(30) Gumbart, J.; Khalili-Araghi, F.; Sotomayor, M.; Roux, B. 2012. Constant electric field simulations of the membrane potential illustrated with simple systems. Biochim. Biophys. Acta, Biomembr. 1818:294–302. doi: 10.1016/j.bbamem.2011.09.030.

(31) Castillo, J. P.; Rui, H.; Basilio, D.; Das, A.; Roux, B.; Latorre, R.; Bezanilla, F.; Holmgren, M. 2015. Mechanism of potassium ion uptake by the Na+/K+-ATPase. Nat. Commun. 6:7622. doi: 10.1038/ncomms8622.

(32) Bignucolo, O.; Chipot, C.; Kellenberger, S.; Roux, B. 2022. The Galvani Offset Potential and Constant-pH Simulations of Membrane Proteins. J. Phys. Chem. B. 126:6868–6877. doi: 10.1021/acs.jpcb.2c04593.

(33) Evans, M. G.; Polanyi, M. 1935. Some applications of the transition state method to the calculation of reaction velocities, especially in solution. Trans. Faraday Soc. 31:875–894. doi: 10.1039/tf9353100875.

(34) Carlsen, R.; Weckel-Dahman, H.; Swanson, J. M. J. 2024. Nernst equilibrium, rectification, and saturation: Insights into ion channel behavior. Biophys. J. 123:4304–4315. doi: 10.1016/j.bpj.2024.10.016.

(35) Nernst, W. 1889. Die elektromotorische wirksamkeit der jonen. Z. Phys. Chem. 4:129–181.

(36) Zakharian, E.; Reusch, R. N. 2004. Streptomyces lividans potassium channel KcsA is regulated by the potassium electrochemical gradient. Biochem. Biophys. Res. Commun. 316:429–436. doi: 10.1016/j.bbrc.2004.02.069.

(37) Nimigean, C. M.; Chappie, J. S.; Miller, C. 2003. Electrostatic Tuning of Ion Conductance in Potassium Channels. Biochemistry. 42:9263–9268. doi: 10.1021/bi0348720.

(38) Allen, T. W.; Kuyucak, S.; Chung, S. H. 1999. Molecular dynamics study of the KcsA potassium channel. Biophys. J. 77:2502–2516. doi: 10.1016/s0006-3495(99)77086-4.

(39) Bernèche, S.; Roux, B. 2000. Molecular dynamics of the KcsA K(+) channel in a bilayer membrane. Biophys. J. 78:2900–2917. doi: 10.1016/s0006-3495(00)76831-7.

(40) Egwolf, B.; Roux, B. 2010. Ion Selectivity of the KcsA Channel: A Perspective from Multi-Ion Free Energy Landscapes. J. Mol. Biol. 401:831–842. doi: 10.1016/j.jmb.2010.07.006.

(41) Kopec, W.; Rothberg, B. S.; de Groot, B. L. 2019. Molecular mechanism of a potassium channel gating through activation gate-selectivity filter coupling. Nat. Commun. 10: 5366. doi: 10.1038/s41467-019-13227-w.

(42) Hui, C.; de Vries, R.; Kopec, W.; de Groot, B. L. 2025. Effective polarization in potassium channel simulations: Ion conductance, occupancy, voltage response, and selectivity. Proc. Natl. Acad. Sci. U. S. A. 122:e2423866122. doi:10.1073/pnas.2423866122.

(43) Furini, S.; Catacuzzeno, L. 2026. Current-voltage characteristics of K+ channels estimated by MD simulations and Markov State Models. bioRxiv. 2026.2002. 2004.703779.

(44) Zhou, Y.; Morais-Cabral, J. H. 2001. Chemistry of ion coordination and hydration revealed by a K+ channel-Fab complex at 2.0 A resolution. Nature. 414:43–48. doi: 10.1038/35102009.

(45) Bhate, M. P.; Wylie, B. J.; Tian, L.; McDermott, A. E. 2010. Conformational dynamics in the selectivity filter of KcsA in response to potassium ion concentration. J. Mol. Biol. 401:155–166. doi: 10.1016/j.jmb.2010.06.031.

(46) Ryan, M. J.; Gao, L.; Valiyaveetil, F. I.; Kananenka, A. A.; Zanni, M. T. 2023. Probing Ion Configurations in the KcsA Selectivity Filter with Single-Isotope Labels and 2D IR Spectroscopy. J. Am. Chem. Soc. 145:18529–18537. doi: 10.1021/jacs.3c05339.

(47) Ryan, M. J.; Gao, L.; Valiyaveetil, F. I.; Kananenka, A. A.; Zanni, M. T. 2024. Water inside the Selectivity Filter of a K+ Ion Channel: Structural Heterogeneity, Picosecond Dynamics and Hydrogen Bonding. J. Am. Chem. Soc. 146:1543–1553. doi: 10.1021/jacs.3c11513.

(48) Bhate, M. P.; McDermott, A. E. 2012. Protonation state of E71 in KcsA and its role for channel collapse and inactivation. Proc. Natl. Acad. Sci. U. S. A. 109:15265–15270. doi:10.1073/pnas.1211900109.

(49) Wylie, B. J.; Bhate, M. P.; McDermott, A. E. 2014. Transmembrane allosteric coupling of the gates in a potassium channel. Proc. Natl. Acad. Sci. U. S. A. 111:185–190. doi:10.1073/pnas.1319577110.

(50) Xu, Y.; Bhate, M. P.; McDermott, A. E. 2017. Transmembrane allosteric energetics characterization for strong coupling between proton and potassium ion binding in the KcsA channel. Proc. Natl. Acad. Sci. U. S. A. 114:8788–8793. doi:10.1073/pnas.1701330114.

(51) Sun, Z.; Xu, Y.; Zhang, D.; McDermott, A. E. 2020. Probing allosteric coupling in a constitutively open mutant of the ion channel KcsA using solid-state NMR. Proc. Natl. Acad. Sci. U. S. A. 117:7171–7175. doi:10.1073/pnas.1908828117.

