## Supplemental Information for "Voltage versus concentration in ion channels: Physical signatures reveal rate-limiting steps"

**Supplemental methods**

**Derivation for voltage-responsive rate constants and biomolecular flux**

In previous works (1-4), we calculated the voltage coupling factor ($\varphi\left( z \right)$) using the linear coupling method (5,6), which takes the difference between ensemble-averaged electrostatic potentials for simulations run with a negative (${\Delta\phi}_{elec}^{-V}(z)$) and positive (${\Delta\phi}_{elec}^{+V}(z)$) applied electric field of equal magnitude and normalizes the expression by the total voltage drop across the simulation box ($V_{T}$):

$$\varphi\left( z \right)= \frac{{\Delta\phi}_{elec}^{-V}(z)-{\Delta\phi}_{elec}^{+V}(z)}{V_{T}}$$

(S1)

In kinetic modeling, rate constants of rare-event transitions depend on the difference in free energy between meta-stable intermediates and transition barriers (${\Delta G}_{ij}^{\ddagger}$). Using Roux’s W-route (7), we previously described the free energy difference under a transmembrane potential as (4):

$$G\left( z,V \right)= \varphi\left( z \right)qV+G(z,0)$$

(S2)

$${\Delta G}_{ij}^{\ddagger}\left( z,V \right)=\left[ \varphi\left( z^{t} \right)qV+G\left( z^{t},0 \right) \right]-\left[ \varphi\left( z^{r} \right)qV+G\left( z^{r},0 \right) \right]$$

(S3)

where q is the charge of the ion moving in the transition, z^t^ and z^r^ are the transition barrier and reactant basin locations along the reaction path, respectively, and $G_{z;0}^{t}$ and $G_{z;0}^{r}$ are the zero-voltage free energy at the transition barrier and reactant basin, respectively. Using the Eyring-Polanyi (8) equation, given by Eq. 4, we can then derive a relationship between rate coefficients under different transmembrane potentials (4):

$$k_{ij}^{0}=\frac{\omega_{0}}{2\pi}e^{\frac{-{\Delta G}_{ij}^{\ddagger}(z, 0)}{k_{B}T}}$$

(S4)

$$k_{ij}(V)=k_{ij}^{0}e^{\frac{-\left[ \varphi_{ij}\left( z^{t} \right)-\varphi_{ij}\left( z^{r} \right) \right]qV}{k_{B}T}}$$

(S5)

where $k_{B}$ is Boltzmann’s constant, T is the temperature (K), $\omega_{0}$ is the attempt frequency, and $k_{ij}(V)$ and $k_{ij}^{0}$ are the rate coefficients for a transition from state i to j with and without the influence of voltage, respectively (4,9).

For voltage responsive rates, the flux through a given transition is:

$$\mathcal{J}_{ij}=P_{i}k_{ij}(V)$$

(S6)

where $\mathcal{J}_{ij}$ is the flux from state i to j and $P_{i}$ the population of state i, which is equivalent to the probability for a single channel. Uptake rate coefficients are modified by the bulk ionic concentration in mM of the ion of interest (${[ion]}_{bulk}):$

$$k_{ij}^{bind}(V)={{[ion]}_{bulk}k}_{ij}^{0}e^{\frac{-\left[ \varphi_{ij}\left( z^{t} \right)-\varphi_{ij}\left( z^{r} \right) \right]qV}{k_{B}T}}$$

(S7)

And the subsequent flux to:

$$\mathcal{J}_{ij}={{[ion]}_{bulk}P}_{i}k_{ij}^{0}e^{\frac{-\left[ \varphi_{ij}\left( z^{t} \right)-\varphi_{ij}\left( z^{r} \right) \right]qV}{k_{B}T}}$$

(S8)

At steady state assumption, the flux moving through one plane dividing binding sites is the net flux ($\mathcal{J}_{Net})$ moving through the protein. We can then calculate $\mathcal{J}_{Net}$ through the protein as:

$$\mathcal{J}_{Net}= \sum_{ij} P_{i}k_{ij}(V)-\sum_{ji} P_{j}k_{ji}(V)$$

(S9)

**Relating electrostatics to rate constants in MsRKM**

Since free energy and electric potential energy are additive, ${\Delta G}^{\ddagger}$ for each transition can be expressed as a summation of the activation energy of the empty channel and the electric potential energy due to electrostatic attractive/repulsive forces between ions.

$${\Delta G}_{ij}^{\ddagger}= {\Delta G}_{empty}^{\ddagger}+ {\Delta E}_{elec}$$

(S10)

The electric potential energy (E_elec_) between two point charges q_m_ and q_n_ separated by a distance r is:

$$E_{elec}= \frac{1}{4\pi\varepsilon_{0}\varepsilon'}\frac{q_{m}q_{n}}{r_{mn}}$$

(S11)

where $\varepsilon_{0}$ is the permittivity of free space and $\varepsilon'$ is the effective dielectric constant. The effective dielectric constant is a singular term that represents both the dielectric constant of the environment as well as any micro-scale shielding effects and minor conformational rearrangements that modify the potential energy (10). The effect of this electric potential energy on rate constants can be calculated for a particular transition as the net difference in electric potential energy between the transition state and reactant states:

$${\Delta E}_{elec}= \frac{1}{4\pi\varepsilon_{0}\varepsilon'}q_{m}q_{n}\left( \frac{1}{r_{mn}^{T}}-\frac{1}{r_{mn}^{R}} \right)$$

(S12)

where $r_{mn}^{T}$ and $r_{mn}^{R}$ are the distance between a transiting ion at the transition state or reactant position, respectively, and an ion at a different binding site within the protein.

Combining Eq. S10 with Eq. S5 we get:

$$k_{ij}(V)= \frac{\omega_{0}}{2\pi}e^{- \frac{{\Delta G}_{empty}^{\ddagger}(V)+{\Delta E}_{elec}}{k_{B}T}}$$

(S13)

$$k_{ij}(V)= {k_{ij}^{Base}(V)e}^{- \frac{{\Delta E}_{elec}}{k_{B}T}}$$

(S14)

where $k_{ij}^{Base}(V)$ is the transition rate from state i to j for a channel or transporter empty of ions other than the transiting ion.

**Voltage coupling profile**

The fraction of the membrane potential, $\varphi(z)$, captures the normalized fraction of the voltage drop across a membrane that an ion would feel as it traverses from one side of the membrane to the other. In previous works, $\varphi(z)$ has been calculated using explicit solvent (7,11-14) or a linearized PB-V continuum approximation (2,4,5). In this paper, all model proteins used the symmetrical coupling factor in Fig. S1 (1,2,4). This approximation models the voltage profile along the z-axis of a simulated protein while removing any protein-specific influences on the voltage.

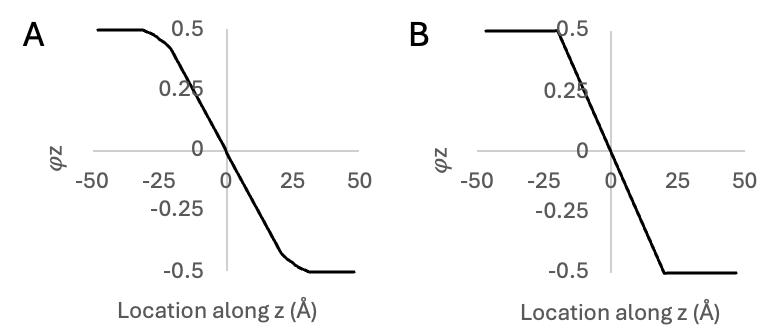

Figure S1 (A) The synthetic voltage coupling profile used in this study is perfectly symmetric about 0.

**Model specifications**

For all models, we used a generic membrane thickness of 40 Å. Unless specified differently in the text below, binding site locations for electrostatic calculations were placed equidistant from each other and the membrane boundary. Transition states (TS) are located at the halfway distance between binding sites or between a binding site and the adjacent membrane boundary unless otherwise specified.

*Perfectly symmetric model*

The two site perfectly symmetric model was designed to maximize the total symmetry in the system. Thus, Δ𝜑 values are equal and $k_{ij}^{Base}$ are equal at intracellular/extracellular K^+^ concentrations of 100 mM. Binding sites were placed at 𝜑 values of ± $\sim0.1\bar{6}$ and binding site were $\sim13.\bar{3}$ Å apart from one another. A constant $\epsilon'$ value of 200 was used for all non-$k_{ij}^{Base}$ transitions. This results in release rate constant increases for the [11] state and uptake rate constants decrease for the [01] and [10] states of ~33%.

*Additional models*

In order to pull trends from changing $k_{ij}$ and Δ𝜑 values, electrostatic effects were kept consistent with the two site perfectly symmetric model. Unless specified otherwise, intracellular and extracellular K^+^ concentrations remained 100 mM. Models are presented in order as discussed in the main text.

|  | Model | ${k_{ij}}^{Uptake}$ | ${k_{ij}}^{Release}$ | ${k_{ij}}^{Transfer}$ | $\vert\Delta{\varphi\vert}^{all}$ |
| --- | --- | --- | --- | --- | --- |
| I-V vs. I-μ Analyses | Fig. 4 | 0.1 | 1 | 1 | $0.1\bar{6}$ |
|  | Fig. 5 | 1 | 0.1 | 1 | $0.1\bar{6}$ |
|  | Fig. 6 | 1 | 1 | 0.1 | $0.1\bar{6}$ |

Table S1: MsRKM model specifications used for all two site mirror symmetric models. $k_{ij}^{X}$ is the relative weighting of the base rate constants. For ${k_{ij}}^{Uptake}$, this relative weighting is calculated at 100 mM extracellular/intracellular K^+^ concentration. ${|\Delta\varphi|}^{all}$ is the absolute $\Delta\varphi$ value used for all transitions.

|  | Model | $k_{ij}^{Bind}$ | $k_{ij}^{Release}$ | $k_{ij}^{Transfer}$ | ${\vert\Delta\varphi\vert}^{Release}$ | ${\vert\Delta\varphi\vert}^{U/T}$ |
| --- | --- | --- | --- | --- | --- | --- |
| Measured vs. Predicted Analyses | Fig. 7 | 1 | 0.01 | 1 | 0.26 | 0.1165 |
|  | Fig. 9 *A* | 0.01 | 1 | 1 | $0.1\bar{6}$ | $0.1\bar{6}$ |
|  | Fig. 9 *B* | 1 | 1 | 0.01 | $0.1\bar{6}$ | $0.1\bar{6}$ |
|  | Fig. 9 *C* | 1 | 0.01 | 1 | $0.1\bar{6}$ | $0.1\bar{6}$ |

Table S2: MsRKM model specifications used for models in the main text. $k_{ij}^{X}$ is the relative weighting of the base rate constants. For ${k_{ij}}^{Uptake}$, this relative weighting is calculated at 100 mM extracellular/intracellular K^+^ concentration. ${|\Delta\varphi|}^{X}$is the absolute $\Delta\varphi$ value used for the indicated transition.

**Supplemental results**

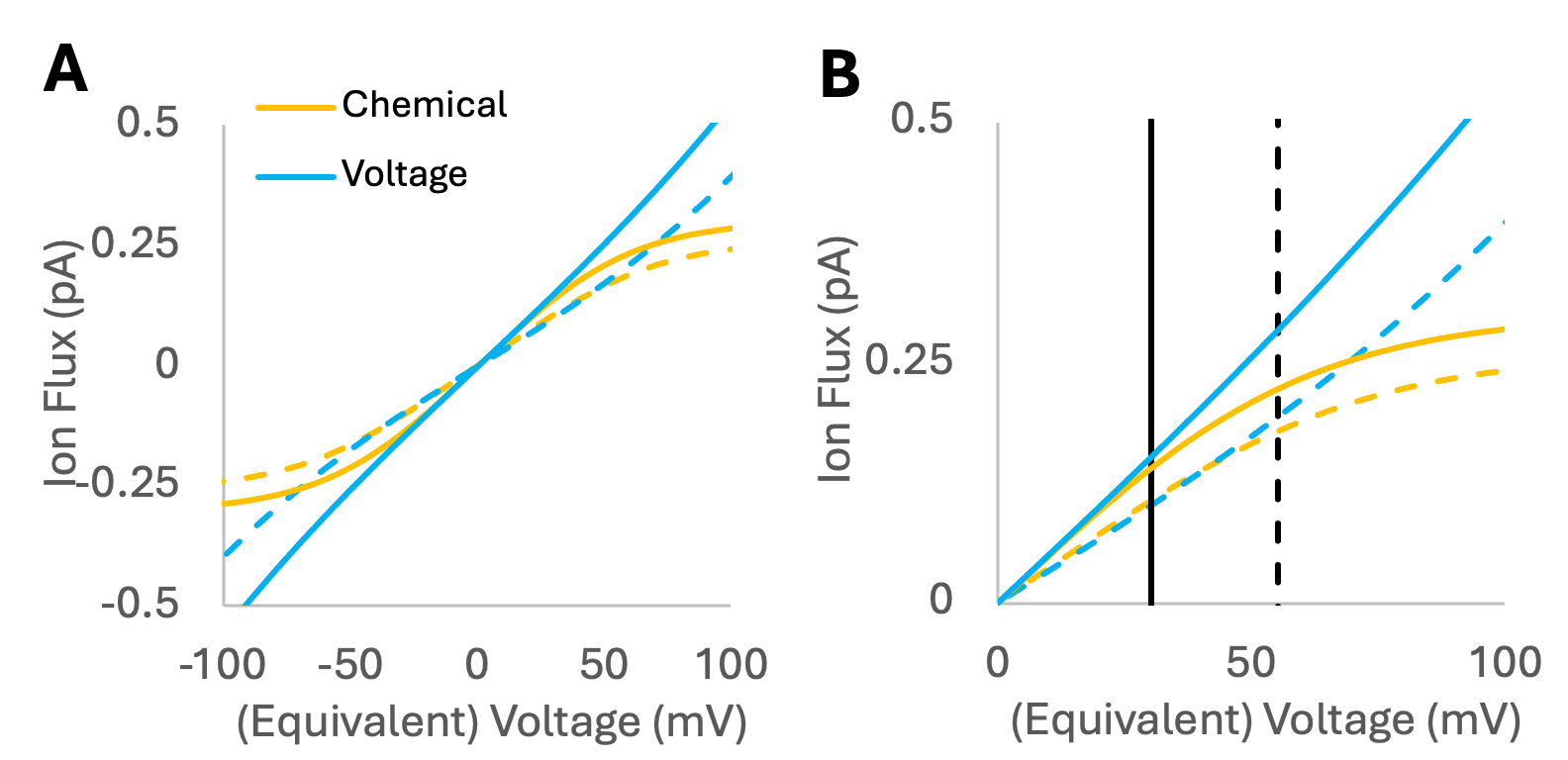

Figure S2 (A) The voltage- (blue) and chemical potential- (yellow) driven flux for two site model systems with a transfer RLS. Solid lines are for a model system where transfer is 10x slower than the other rates. Dashed lines are for a model system where transfer is 100x slower than the other rates. (B) Focusing on outward flux, the point at which voltage-driven flux exceeds chemical potential-driven flux (vertical lines) is ~25 mV higher for the model where transfer is more rate limiting.

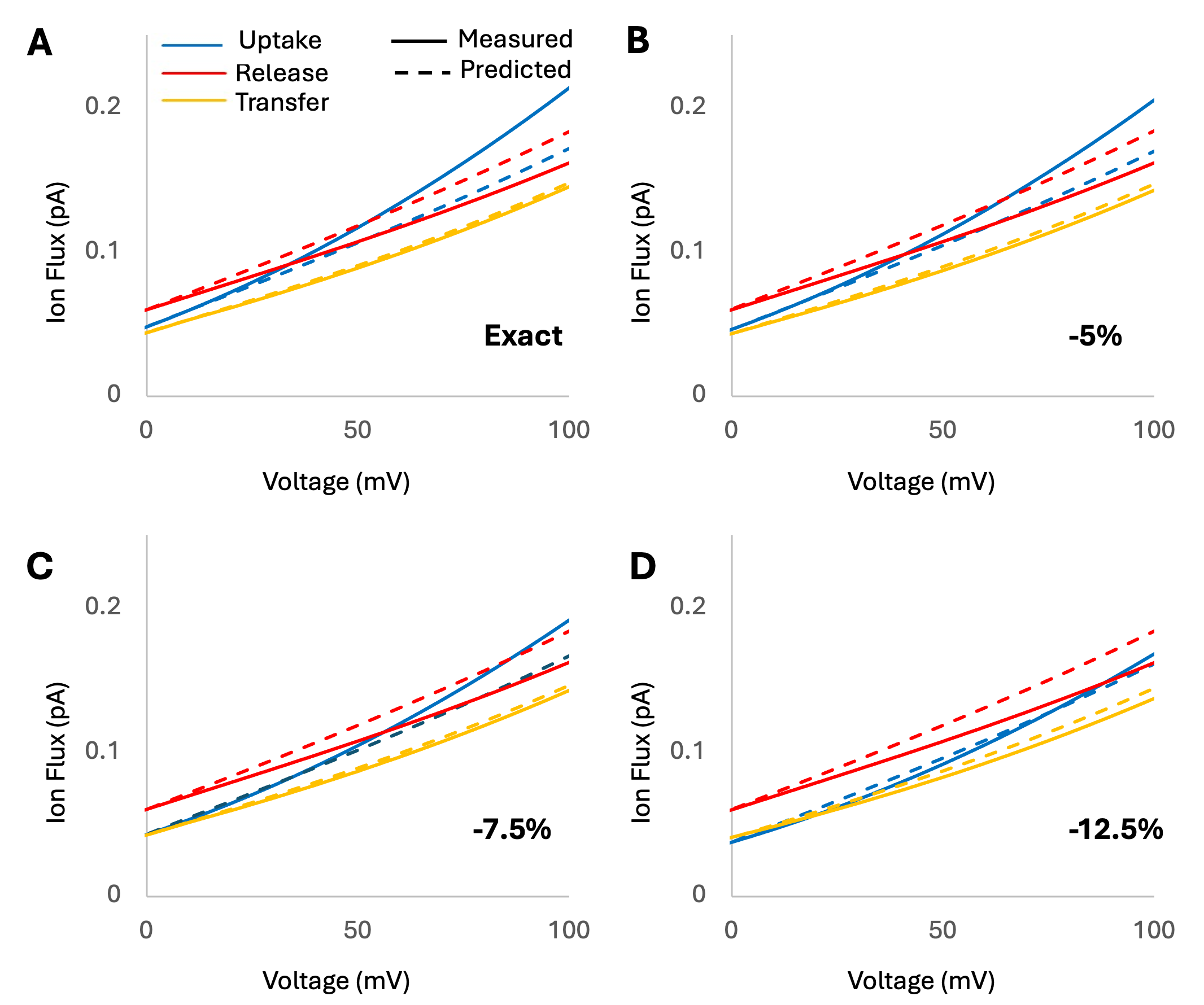

Figure S3 Accuracy of the Predicted vs. Measured assay when net concentrations are not equal. The symmetric net concentration was 200 mM for all assays. The asymmetric net concentration was (A) 200 mM, (B) 190 mM, (C) 185 mM, or (D) 175 mM. The same asymmetric gradient of 1:10 was used in for each assay (~59 mV equivalent). Reliability is lost at 175 mM net concentration (a 12.5% difference from 200 mM), when the uptake-limited system looks transfer-limited, and the transfer-limited system looks uptake-limited .

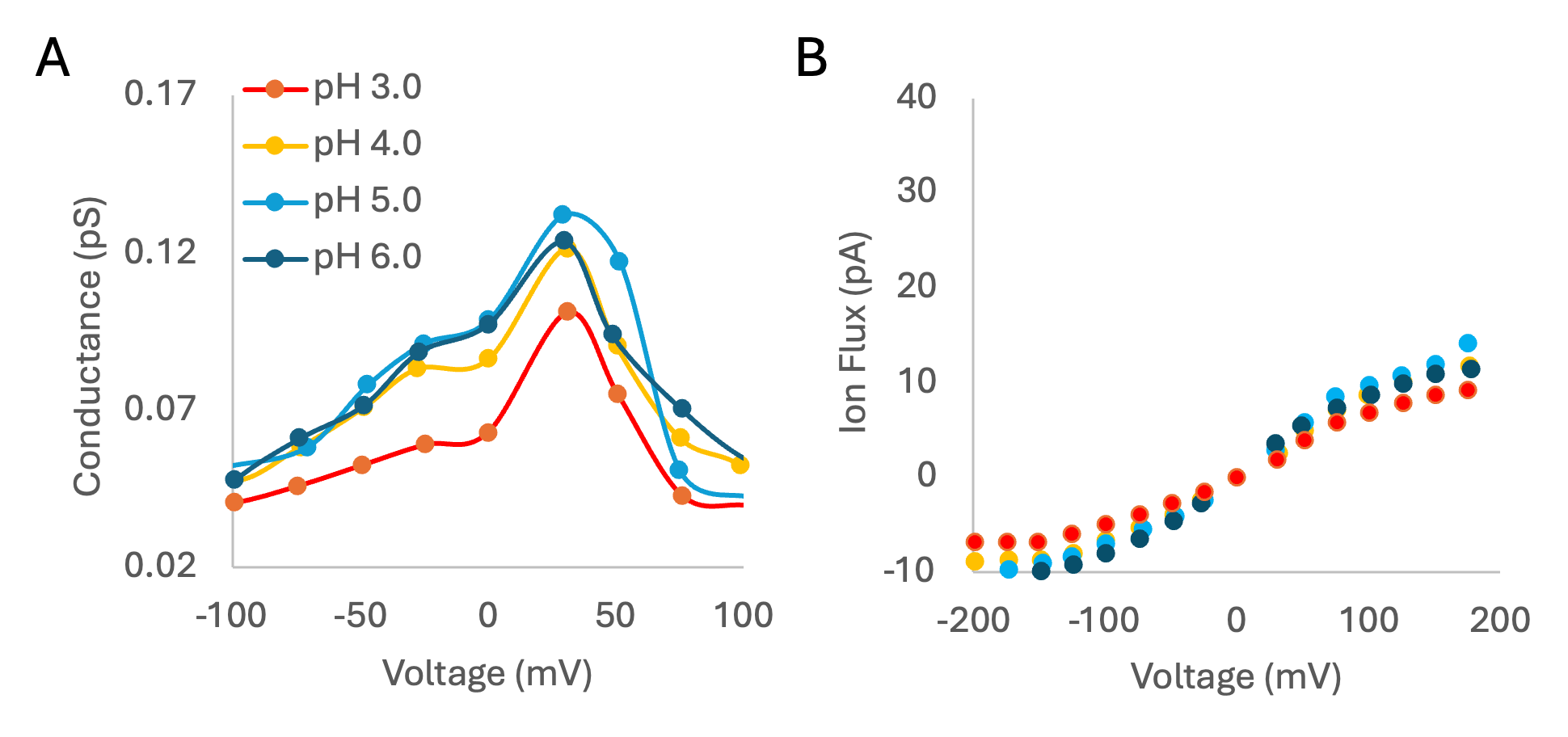

Figure S4 Figure adapted from Nimigean et al. (15): (A) The conductance shape of wild type KcsA extracted from the experimental I-V curves. (B) Potassium flux as a function of pH. Each datapoint represents the average ± SE from at least three separate bilayers.

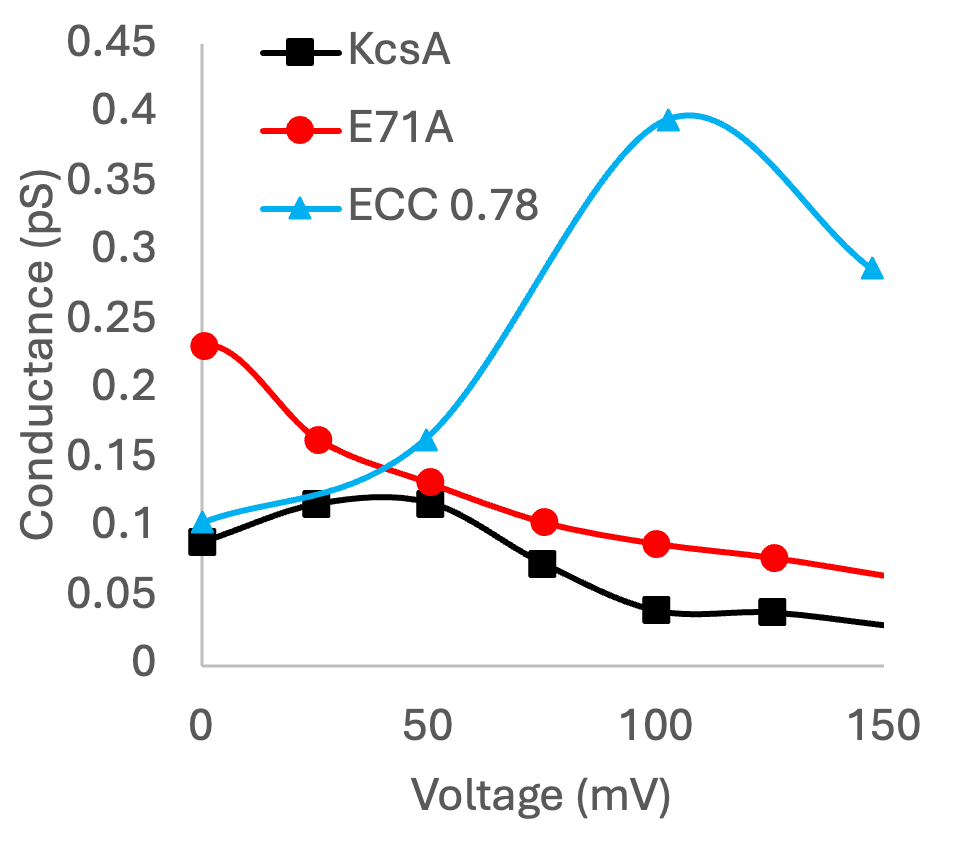

Figure S5 Conductance shapes for wild type KcsA (black squares) and E71A mutant (red circles) from I-V curves with symmetric 100 mM K^+^ concentration as taken from Cheng et al. (16) Additional simulation data provided from Hui et al. (17) at symmetric 500 mM K^+^ concentration (blue triangles).

**References**

(1) Carlsen, R.; Weckel-Dahman, H.; Swanson, J. M. J. 2024. Nernst equilibrium, rectification, and saturation: Insights into ion channel behavior. *Biophys. J.* 123:4304-4315. doi: 10.1016/j.bpj.2024.10.016.

(2) Weckel-Dahman, H.; Carlsen, R.; Daum, A.; He, M.; Southam, T. G.; Swanson, J. M. J. 2025. Ion Channel Reaction Networks: Dielectric Screening and the Importance of Off-Pathway Flux. *J. Chem. Theory Comp.* 21:3700-3711. doi: 10.1021/acs.jctc.4c01569.

(3) Weckel-Dahman, H.; Carlsen, R.; Daum, A.; Swanson, J. M. J. 2026. From Flux to Function: Extracting Mechanistic Insights from Ion Channels via I–V and I–μ Analyses. *J. Phys. Chem. B.* 130:2733-2742. doi: 10.1021/acs.jpcb.5c06019.

(4) Weckel-Dahman, H.; Carlsen, R.; Swanson, J. M. J. 2024. Multiscale Responsive Kinetic Modeling: Quantifying Biomolecular Reaction Flux under Varying Electrochemical Conditions. *J. Chem. Theory Comp.* 21:374-389. doi: 10.1021/acs.jctc.4c00872.

(5) Castillo, J. P.; Rui, H.; Basilio, D.; Das, A.; Roux, B.; Latorre, R.; Bezanilla, F.; Holmgren, M. 2015. Mechanism of potassium ion uptake by the Na+/K+-ATPase. *Nat. Commun.* 6:7622. doi: 10.1038/ncomms8622.

(6) Gumbart, J.; Khalili-Araghi, F.; Sotomayor, M.; Roux, B. 2012. Constant electric field simulations of the membrane potential illustrated with simple systems. *Biochim. Biophys. Acta, Biomembr.* 1818:294-302. doi: https://doi.org/10.1016/j.bbamem.2011.09.030.

(7) Roux, B. 2008. The Membrane Potential and its Representation by a Constant Electric Field in Computer Simulations. *Biophys. J.* 95:4205-4216. doi: 10.1529/biophysj.108.136499.

(8) Eyring, H. 2002. The Activated Complex and the Absolute Rate of Chemical Reactions. *Chem. Rev*. 17:65-77. doi: 10.1021/cr60056a006.

(9) Mayes, H. B.; Lee, S.; White, A. D.; Voth, G. A.; Swanson, J. M. J. 2018. Multiscale Kinetic Modeling Reveals an Ensemble of Cl–/H+ Exchange Pathways in ClC-ec1 Antiporter. *J. Amer. Chem. Soc.* 140:1793-1804. doi: 10.1021/jacs.7b11463.

(10) Sieradzan, A. K.; Lipska, A. G.; Lubecka, E. A. 2018. Shielding effect in protein folding. *J Mol. Graph Model.* 79*:*118-132. doi: 10.1016/j.jmgm.2017.10.018.

(11) Berneche, S.; Roux, B. 2001. Energetics of ion conduction through the K+ channel. *Nature.* 414:73-77. doi: 10.1038/35102067.

(12) Castillo, J. P.; Rui, H.; Basilio, D.; Das, A.; Roux, B.; Latorre, R.; Bezanilla, F.; Holmgren, M. 2015. Mechanism of potassium ion uptake by the Na+/K+ ATPase. *Nat. Commun.* 6. doi: 10.1038/ncomms8622.

(13) Weckel-Dahman, H.; Carlsen, R.; Swanson, J. M. J. 2024. Multiscale Responsive Kinetic Modeling: Quantifying Biomolecular Reaction Flux under Varying Electrochemical Conditions. *J. Chem. Theory. Comput.* 21:374-389. doi: 10.1021/acs.jctc.4c00872

(14) Bignucolo, O.; Chipot, C.; Kellenberger, S.; Roux, B. 2022. The Galvani Offset Potential and Constant-pH Simulations of Membrane Proteins. *J. Phys. Chem. B.* 126:6868-6877. doi: 10.1021/acs.jpcb.2c04593.

(15) Nimigean, C. M.; Chappie, J. S.; Miller, C. 2003. Electrostatic Tuning of Ion Conductance in Potassium Channels. *Biochemistry*. 42:9263-9268. doi: 10.1021/bi0348720.

(16) Cheng, W. W. L.; McCoy, J. G.; Thompson, A. N.; Nichols, C. G.; Nimigean, C. M. 2011. Mechanism for selectivity-inactivation coupling in KcsA potassium channels. *Proc. Natl. Acad. Sci. U. S. A.* 108:5272-5277. doi:10.1073/pnas.1014186108.

(17) Hui, C.; de Vries, R.; Kopec, W.; de Groot, B. L. 2025. Effective polarization in potassium channel simulations: Ion conductance, occupancy, voltage response, and selectivity. *Proc. Natl. Acad. Sci. U. S. A.* 122:e2423866122. doi:10.1073/pnas.2423866122.
